# Human intracranial correlates of dynamic coding in auditory working memory

**DOI:** 10.1101/2023.08.04.552073

**Authors:** Işıl Uluç, Angelique C. Paulk, Alan Bush, Valentina Gumenyuk, Noam Peled, Parker Kotlarz, Kaisu Lankinen, Fahimeh Mamashli, Nao Matsuda, G. Rees Cosgrove, R. Mark Richardson, Steven M. Stufflebeam, Sydney S. Cash, Jyrki Ahveninen

## Abstract

Working memory (WM), the human brain’s system for maintaining and manipulating information over short timescales, is critical for goal-directed behavior. Classic neurophysiological models, which emphasize persistent delay activity, are being challenged by an idea that WM is maintained through dynamic and activity-silent mechanisms, where the memory trace is preserved by transient changes in the connections between neurons. To date, human evidence for this alternative model has been limited to non-invasive studies. Here, leveraging high-resolution intracranial stereo-EEG with advanced machine learning decoding techniques, we investigate neuronal mechanisms of human auditory WM. Neuronal activity is quantified as broadband high frequency activity (70-190 Hz), a correlate of neuronal firing activity. Our multivariate pattern analyses, validated via robust non-parametric permutation testing, show that WM content is represented in multiple brain regions, exhibiting dynamic rather than persistent delay activity. Crucially, by using task-irrelevant sounds to probe latent mnemonic states, we provide the first human intracranial evidence of activity silent WM maintenance in a local sensory network.

## Introduction

Working memory (WM) is a function crucial for cognition, dynamically storing the range of thoughts required for complex actions. The neuronal mechanisms of WM have been a focus of intensive neuroscience research since the 1970’s (Baddeley and Hitch, 1974; D’Esposito and Postle, 2015). Non-human primate (NHP) and non-invasive human imaging studies have given valuable insight into the brain networks underlying WM processing (Barak & Tsodyks, 2014; Christophel, Klink, Spitzer, Roelfsema, & Haynes, 2017; Constantinidis et al., 2018; D’Esposito & Postle, 2015; Lundqvist, Herman, & Miller, 2018). The emphasis of this research has been shifting from the “where” in the brain is WM maintained to the neurophysiological question of “how” neurons maintain information of WM content (Barak & Tsodyks, 2014; Christophel et al., 2017; D’Esposito & Postle, 2015; Miller, Lundqvist, & Bastos, 2018; Serences, 2016; Stokes, 2015).

The debate surrounding the nature of neuronal processes underlying WM maintenance started with initial WM studies in NHPs, showing that consistent firing in prefrontal cortex (PFC) accompanies WM retention, later on defined as delay activity (Fuster and Alexander, 1971; Rosenkilde et al., 1981). Neurophysiological studies suggested that such sustained delay activity patterns maintain information throughout the WM maintenance period (Goldman-Rakic, 1995; Romo et al., 1999; Constantinidis and Procyk, 2004; Vergara et al., 2016). The sustained firing was shown to be related to both excitatory and inhibitory dynamics of neurons (Barak & Tsodyks, 2014; Sreenivasan & D’Esposito, 2019). Subsequent human neuroimaging studies (Christophel et al., 2012; Christophel and Haynes, 2014; D’Esposito and Postle, 2015), however, showed that content-specific information could also be found in sensory brain areas where no sustained activity is present during WM maintenance. Evidence is also emerging that not only are posterior and sensory areas involved in WM maintenance but also the neuronal processes maintaining WM do not necessarily depend on persistently elevated neuronal firing.

Several more recent neurophysiological studies support an interpretation that persistent firing during the WM delay period is a correlate of other brain functions including attention, decision-making, learning, or motor preparation (Lundqvist et al., 2018a; Sreenivasan and D’Esposito, 2019). At the same time, while the classic studies were based on trial-averaged signals, more recent studies using single-trial analysis approaches (Shafi et al., 2007; Lundqvist et al., 2016) suggest that WM is maintained via more dynamic process, involving sparse (rather than persistent) firing patterns and bursts of neuronal oscillations (Lundqvist et al., 2016; Kucewicz et al., 2017; Bastos et al., 2018; Lundqvist et al., 2018b). During the period between these bursts of activity, WM could be maintained in an “activity-silent” state, which means that the memory trace is preserved by transient changes in the connections between neurons (i.e., short-term synaptic plasticity) that can later reactivate the stored information when needed (Mongillo, Barak, & Tsodyks, 2008; Stokes, 2015) as opposed to persistent activity where population of neurons persistently fire during the memory maintenance period (Goldman-Rakic, 1995; Romo et al., 1999; Constantinidis and Procyk, 2004; Vergara et al., 2016). These transient connectivity changes arise from recent co-activation of neurons during encoding, which transiently strengthens synaptic pathways representing specific content. After firing subsides, these modified connections bias subsequent network dynamics, allowing stored information to be reactivated by transient activation patterns, such as impulse stimuli. However, these alternative theories have been criticized to be inconsistent with many neurophysiological findings regarding the neuronal underpinnings of WM (Constantinidis et al., 2018). For example, a limitation of the activity-silent models is that much of the experimental evidence is coming from non-invasive human studies (Constantinidis et al., 2018).

The insight about WM-related local neuronal circuits that support the persistent firing theory mostly come from NHP studies, primarily conducted in visual and tactile domains (Goldman-Rakic, 1995; Romo et al., 1999; Barak et al., 2010). Yet, using simple delayed-match-to-sample designs in highly trained laboratory animals might make it difficult to differentiate WM processes from other cognitive functions activated in the process, such as adaptive learning and long-term memory. This difficulty is, perhaps, most evident in the domain of auditory WM, as complex auditory-cognitive tasks are very difficult to learn for NHPs (Fritz et al., 2005; Scott et al., 2012; Scott and Mishkin, 2016). Up to this point, this limitation has left us mostly dependent on non-invasive human neuroimaging studies in the auditory domain (Lutzenberger et al., 2002; Kaiser et al., 2007; Kaiser, 2015; Kumar et al., 2016; Uluç et al., 2018; Ahveninen et al., 2023). One way to bridge the gap between NHP neurophysiological models and human neuroimaging studies is to use intracranial EEG recordings in human participants with epilepsy during their presurgical monitoring (Cash & Hochberg, 2015; Parvizi & Kastner, 2018). Utilizing their high signal-to-noise ratio (SNR), human intracranial EEG studies provide direct access to neural correlates of WM compared to non-invasive neuroimaging or magneto-/electroencephalography (MEG/EEG) recordings.

A significant advantage intracranial recordings is that they provide clear access to > 70 Hz broadband high-frequency activity (HFA), which has been suggested to be a close correlate of local neuronal spiking (Ray et al., 2008; Leszczynski et al., 2020). Ray and colleagues (2008) showed a high temporal correlation between HFA and average firing rate in microelectrode recordings, suggesting that HFA is a correlate of multiunit activity (MUA) (although see also Leszczynski et al. 2020). For example, these studies suggest that the power of intracranial activity in the 65-120 Hz range correlates with successful WM performance (Yamamoto et al., 2014). Recent studies suggest that sustained patterns of HFA could play a role also in human WM processing, in line with the persistent firing model (Constantinidis et al., 2018). An open question, therefore, remains whether such WM-related HFA increases shown in humans reflect maintenance or executive processes aiding WM, such as attention.

To dissect the neuronal mechanisms underpinning WM maintenance in humans, we implemented a rigorously controlled retro-cue paradigm and recorded neurophysiological signals from stereo-electroencephalography (sEEG) electrodes during auditory WM maintenance. Auditory tasks were utilized, not only because there are few neurophysiological studies relative to visual and tactile domains but also because auditory cortex is usually well sampled in sEEG implantations due to clinical considerations (Mercier et al., 2022). In our task, we employed carefully designed, nonconceptual complex auditory modulation patterns (ripple sounds) to ensure that the information maintained is nonverbal but has complex spectrotemporal features (for a more detailed explanation, see Mamashli et al., 2021). We then decoded activity patterns from local field potential HFA patterns. We also calculated the averaged HFA to compare the decoding results to the elevated HFA, which is a correlate of multiunit activity (Ray, Crone, Niebur, Franaszczuk, & Hsiao, 2008). We further analyzed the relationship between the response to an unrelated impulse sound with the increase in decoding accuracy. Our results suggest that auditory WM is subserved by a widely distributed and dynamically interacting network of frontoparietal, medial temporal, and sensory regions, which employ a multilayered maintenance mechanism with interacting active and activity silent processes.

## Results

Our multivariate pattern analysis (MVPA) revealed a significant increase in WM decoding accuracy following task-irrelevant impulse sounds in transverse and superior temporal channels encompassing early auditory cortex (AC). Beyond the early AC, WM representations were reliably decoded from dynamic HFA patterns, both during periods of elevated activation and when activity levels did not exceed the pre-trial baseline. **Figure 1** shows the MVPA and HFA analysis pipeline that yielded main neuronal results.

**Figure 1.**
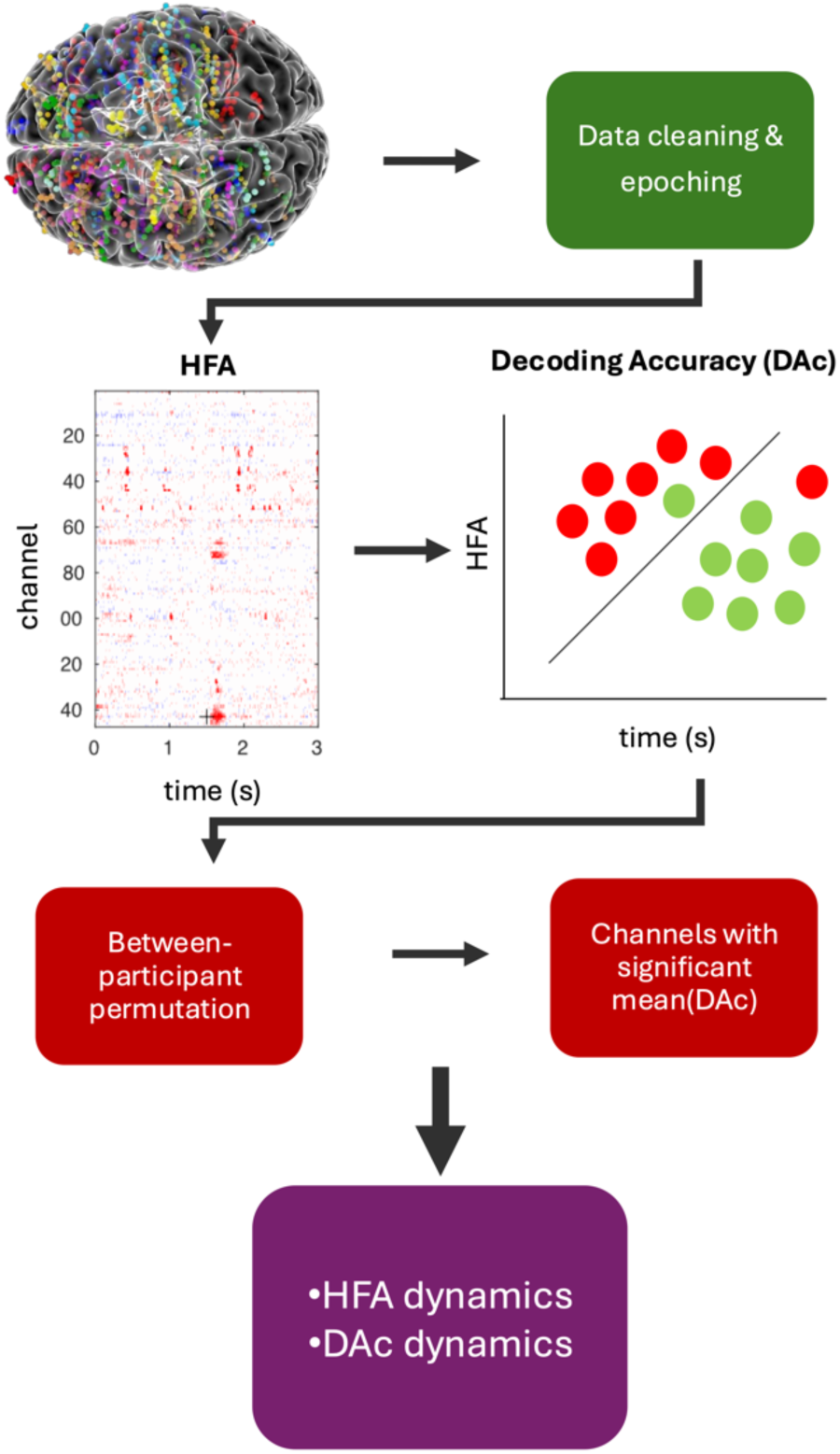
HFA and MVPA analysis pipeline. The data was cleaned and channels showing epileptiform activity were rejected (Janca et al., 2015). Then HFA between 70Hz and 150Hz were averaged and fed into the classifier to obtain decoding accuracies. A between-participant maximum statistics were calculated with 500 permutations. Channels with significant decoding accuracies were selected for further examination for the dynamics of HFA.

### Behavioral Performance

During SEEG, the participants performed a retro-cue WM task with ripple sounds as memoranda. They were asked to ignore the impulse sounds presented 1.5 s into the 3 s maintenance period. In this task, participants (n=9) achieved 67.5±3.4% (mean±SEM) accuracy, substantially above empirical chance responses across all trials of a retro-cue paradigm. We performed an ANOVA with four levels to test if recall performance changes in relation to memorized ripple velocity. Our analysis showed no significant difference between the behavioral performance per memorized ripple velocity. Reaction time analysis could not be calculated due to the self-paced nature of the WM task.

We calculated their behavioral percent correct responses using trials in which the probe either matched the retro-cued item or did not match either the retro-cued or uncued item. Their behavioral performance was significantly higher (p=0.001) in this condition at 77.4±2.8 (mean±SEM).

### Activity silent memory traces in AC: MVPA in AC sites with the strongest response to auditory impulse stimulus

In previous human MEG and EEG studies, activity-silent WM states have been probed by presenting a task-irrelevant impulse stimulus in the middle of the maintenance period (Mamashli et al., 2021; Rose et al., 2016; Uluc et al., 2025; Wolff et al., 2015; Wolff et al., 2017b; Wolff et al., 2020). To provide the first local electrophysiological test in humans from intracranial recordings, we tested in AC channels, whether WM content becomes more readable from the response to an irrelevant impulse sound. To this end, this analysis was focused on participants with contacts in early AC, in the channels that in each participant showed the largest early HFA increase to these impulse sounds. If the WM content were carried in activity-silent states, such an impulse sound should help reveal those silent states from the evoked response (Mamashli et al., 2021; Rose et al., 2016; Uluc et al., 2025; Wolff et al., 2015; Wolff et al., 2017b; Wolff et al., 2020). At the group level, we observed a statistically significant increase in WM content decoding accuracy following the impulse sound. **Figure 2** shows the group median time courses for the impulse-evoked HFA response and the MVPA decoding accuracy in the analyzed auditory cortex channels. This constitutes compelling evidence for activity-silent WM maintenance in early AC.

**Figure 2.**
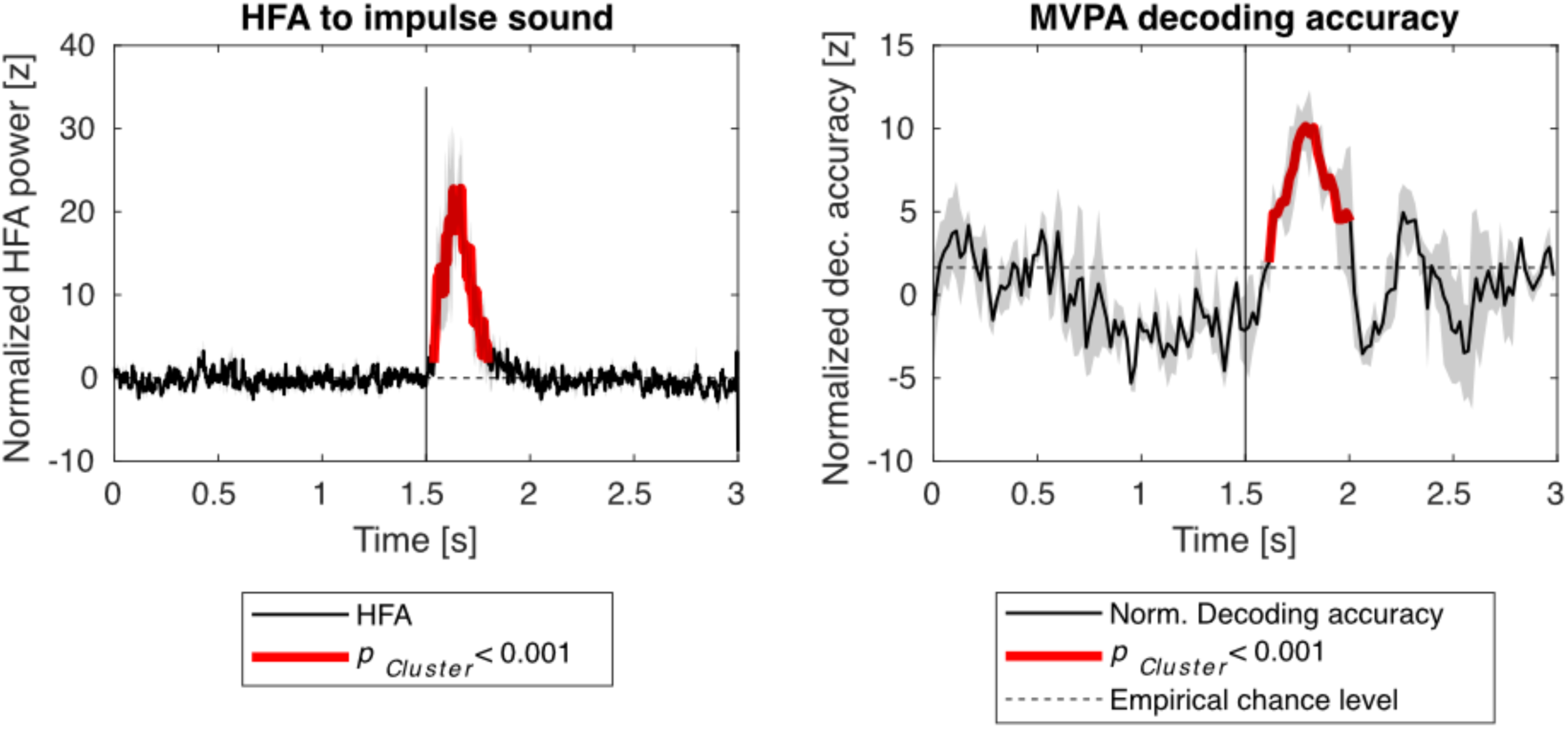
Enhancement of MVPA decoding accuracy after the impulse sound. Group median values from five participants are shown, from each participant’s contact showing the strongest early HFA power after the impulse sound onset in auditory cortex. The HFA power values have been normalized relative to the pre-impulse baseline; the decoding accuracies based on each participant’s permutation distribution. The group statistics are based on a non-parametric cluster-based randomization test (N=5).

These findings are also evident at an individual level. **Figure 3** shows HFA responses to the impulse sound and the corresponding decoding accuracy time course for two representative participants. The decoding accuracy increases above empirical chance level for both channels after the impulse sound with the HFA increase in response to the impulse sound.

**Figure 3.**
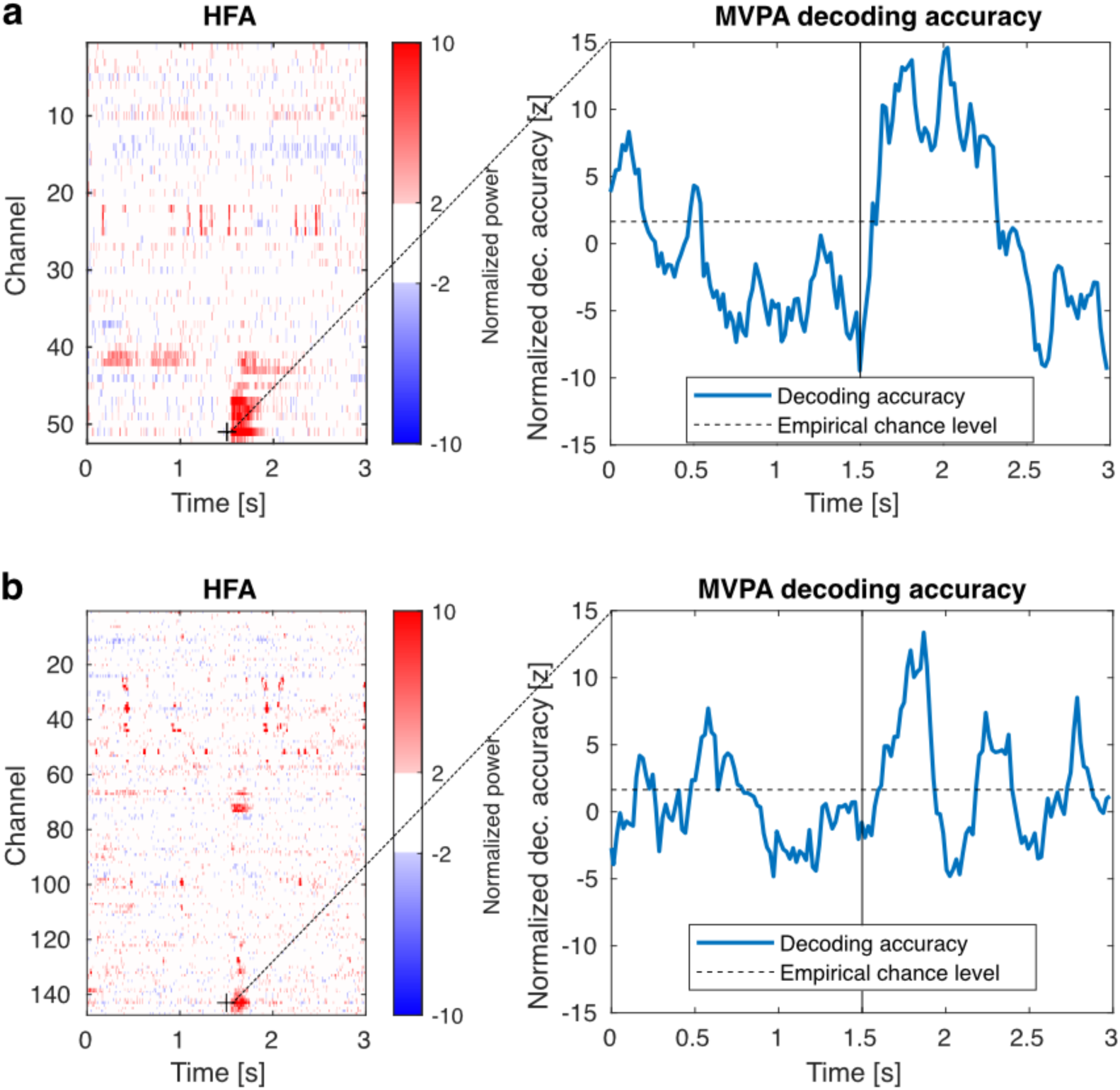
Enhancement of MVPA decoding accuracy after the impulse sound in two participants (N=2; a-b). The HFA power values from all channels are shown at the left. The MVPA decoding accuracy from the contact showing the strongest early-latency HFA after the impulse sound is shown on the right (marked with a cross on the left panel). Empirical chance level has been defined based on a permutation distribution (p<0.05, i.e., z=1.65).

### Multivariate pattern analysis of WM content from HFA

Beyond the MVPA of auditory cortex (AC) channels, we performed a four-class SVM classification on high-frequency activity (HFA) across all channels to probe persistent firing versus dynamic coding at the local level. HFA was used as a proxy for multiunit neuronal firing. We rigorously tested the statistical significance of decoding accuracy using robust non-parametric maximum-statistic controls based on the average of the entire maintenance period, to determine the cortical sites where there was most consistent evidence for WM content maintenance.

Across all participants, our results showed that WM content was widely distributed throughout the brain (**Figure 4**, **Supplementary Figures S2-S9**). **Figure 4a** illustrates the anatomical distribution of significant average decoding accuracies in all channels, as well as the time courses of significant decoding results in the sites for a representative participant, in channels where the average decoding accuracy was above chance level. To examine the temporal evolution of WM representation strength, we also used a cluster-based permutation test to determine statistically significant continuous clusters having better decoding accuracy than empirical chance level (**Figure 4b**). Instead of sustained representations, we found a patchy pattern of clusters of significant decoding accuracy, which are consistent with a dynamic WM maintenance model. These clusters of significant decoding accuracy were found in areas that are related to auditory processing as well as other areas.

**Figure 4.**
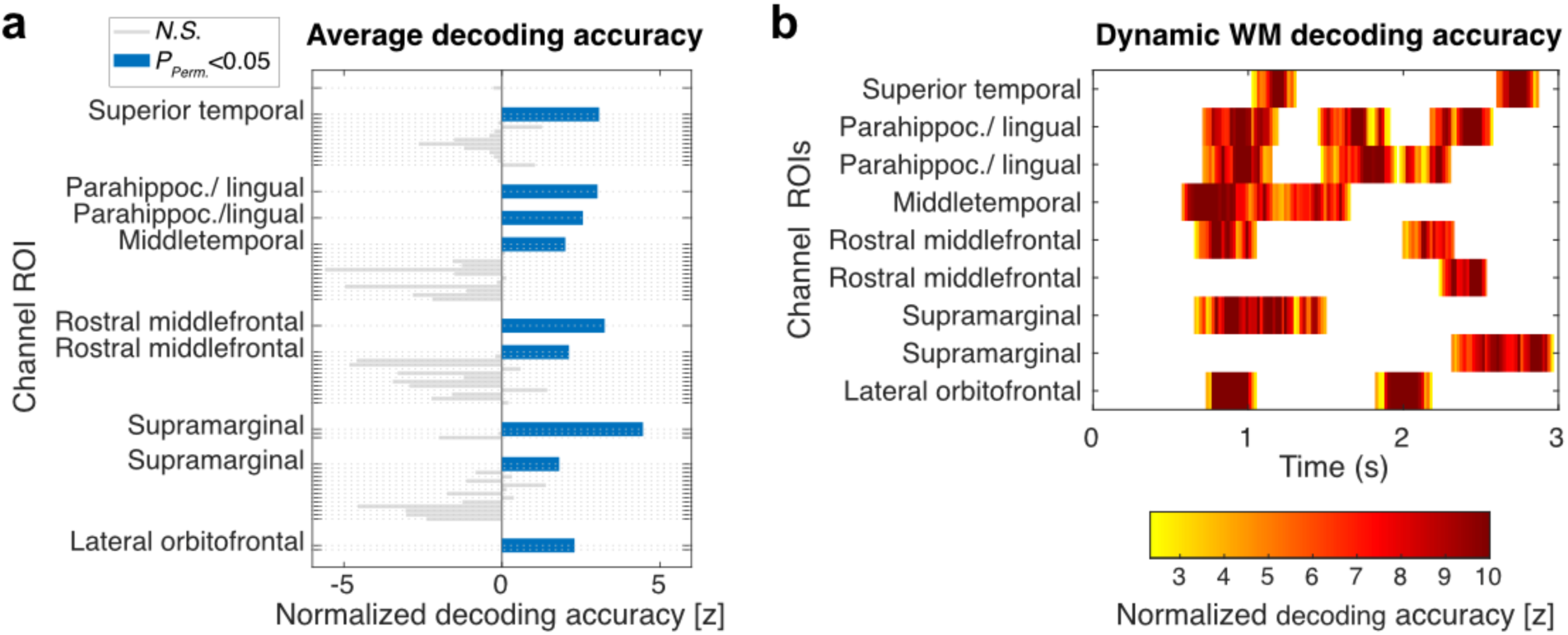
Maintenance period 4-class SVM classification results in a representative participant (N=1), with a high behavioral accuracy in the WM task. (**a)** Average normalized decoding accuracy in all channels. The channels, where decoding accuracy was, on average, significantly above chance level, have been highlighted with thicker, blue bars. Statistical significance was determined by a permutation test with a between-participant maximum-statistic null distribution. All channels were in the right hemisphere in the example participant. (**b)** Time courses for decoding accuracy in channels identified by the cluster-based analysis, focused on those also significant in the time-averaged test. The channels were labeled with their location as it has been determined according to the Freesurfer Desikan atlas. Statistical significance was determined using cluster-based permutation test with a between-participant maximum-statistic null distribution

### Comparing the dynamics of significant decoding accuracy and high-frequency activity

We examined the HFA power for the channels within areas that showed significant decoding accuracy to investigate whether we could find any persistent or elevated HFA activity in the channels that contained WM content information during maintenance based on the average decoding accuracy analysis.

Across participants, HFA power during the trial exhibited highly dynamic, non-persistent temporal profiles in channels carrying WM-related information. **Figure 5** shows HFA power across the whole trial in a representative participant, in the channels where the average decoding accuracy across the maintenance period was statistically significant. In this participant, the HFA time courses were highly dynamic and variable in channels that carried information of WM content. For example, in the two middle frontal channels with a high average decoding accuracy in this participant, the elevated maintenance related HFA was restricted to the periods immediately following the retro-cue, which may reflect attention related responses. In these same channels, HFA was also elevated in response to stimuli during encoding and recall periods. Similar dynamic HFA time courses were observed across all participants (see **Supplementary Figures S10–S17** for results in remaining participants). Notably, no channels exhibited persistent HFA enhancement; if anything, the only sustained effects were suppressive. In particular, two channels from participant 9 where persistent HFA suppression has been found in left pars triangularis and highly dense HFA suppression in right temporal pole and right medial orbitofrontal channels.

**Figure 5.**
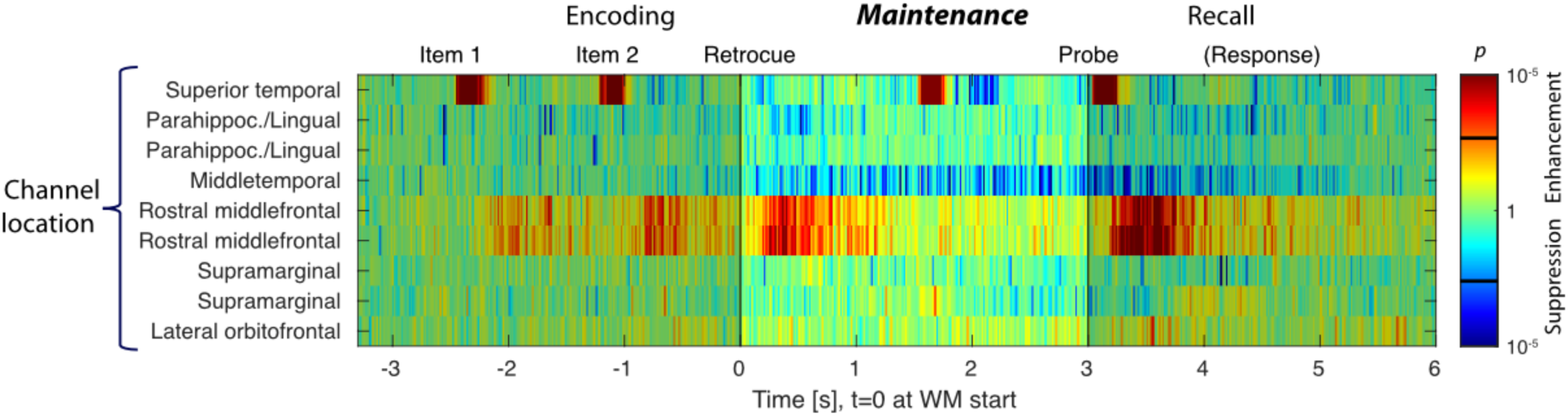
Whole-trial averaged HFA analysis from the channels that carried information of the WM content in a participant with high behavioral accuracy (N=1). Difference relative to the pre-trial baseline is shown. The black bars over the color bar denote the critical *p* value determined using the FDR procedure (Benjamini & Hochberg, 1995). The time periods before and after WM maintenance period are shaded. All channels were in the right hemisphere in the example participant.

The relationship between HFA dynamics and MVPA decoding during the maintenance period was highly consistent across participants (**Figure 6**; **Supplementary Figures S18–S25**). **Figure 6** shows a side-by-side comparison of HFA and MVPA results for the maintenance period in the same participant as in **Figure 5**: Continuous clusters of significant decoding were found in frontal, parahippocampal, and temporal channels. The significant decoding periods in these channels did not persist throughout the whole maintenance period but fluctuated over time, appearing and disappearing intermittently. HFA analysis on the same channels revealed intermittent bursts of activity, consistent with transient coding states rather than evidence for persistent increases. This HFA vs. MVPA pattern was observed in all participants. The only exception was Participant 9 where there was evidence of persistent HFA suppression in pars triangularis, temporal pole and medial orbitofrontal channels. Taken together, our HFA and MVPA results attest to the multi-site, dynamic nature of WM maintenance. Interestingly, periodic suppression was observed in several temporal channels during the maintenance phase, which may constitute a neurophysiological signature of synaptic short term plasticity (Barak & Tsodyks, 2014). According to previous studies, such synaptic facilitation leads to transient bursts of enhanced activity driven by accumulating synaptic efficacy, typically followed by a suppression phase that prevents runaway excitation. This periodic suppression may serve as a dynamic signature of underlying synaptic facilitation mechanisms in recurrent neural circuits (Barak & Tsodyks, 2014).

**Figure 6.**
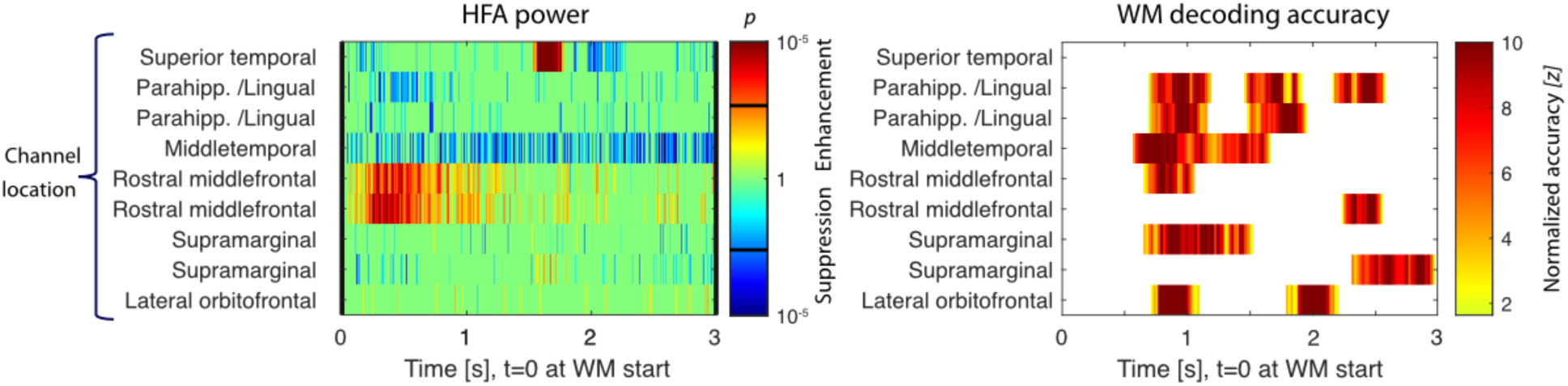
Comparison of averaged HFA analysis and MVPA results for WM maintenance period (N=1). The data of a representative participant with a high behavioral accuracy have been shown. (Left) HFA power during the maintenance period. Difference relative to the pre-trial baseline is shown. The black bars over the color bar denote the critical p value determined as FDR (Benjamini & Hochberg, 1995). (Right) MVPA time courses. The decoding results have been thresholded based on a cluster-based permutations test with a between-participant maximum-statistic null distribution.

### Group level anatomical distribution of channels with elevated HFA and decoding

To further investigate potential elevated activity that may not have been captured by dynamic estimates, we employed a more sensitive temporal averaging approach by calculating the mean HFA over the maintenance period in group level. This analysis identified a subset of channels where average HFA was significantly higher during maintenance compared to the pre-trial baseline. We then compared the number of these channels showing elevated activity with those exhibiting significant decoding in time-averaged estimates during the maintenance period. **Figure 7** shows the distribution of channels with elevated HFA and significant WM decoding, organized by anatomical parcels of the Freesurfer Desikan atlas (Desikan et al., 2006). While areas like the rostral middle frontal and middle temporal cortex included channels with both elevated HFA and significant decoding, other areas show significant decoding without elevated HFA. These findings indicate that although sustained increases in HFA amplitude are present in some regions during WM maintenance, decodable information can also be carried by temporally structured patterns of activity that do not manifest as elevated mean HFA.

**Figure 7.**
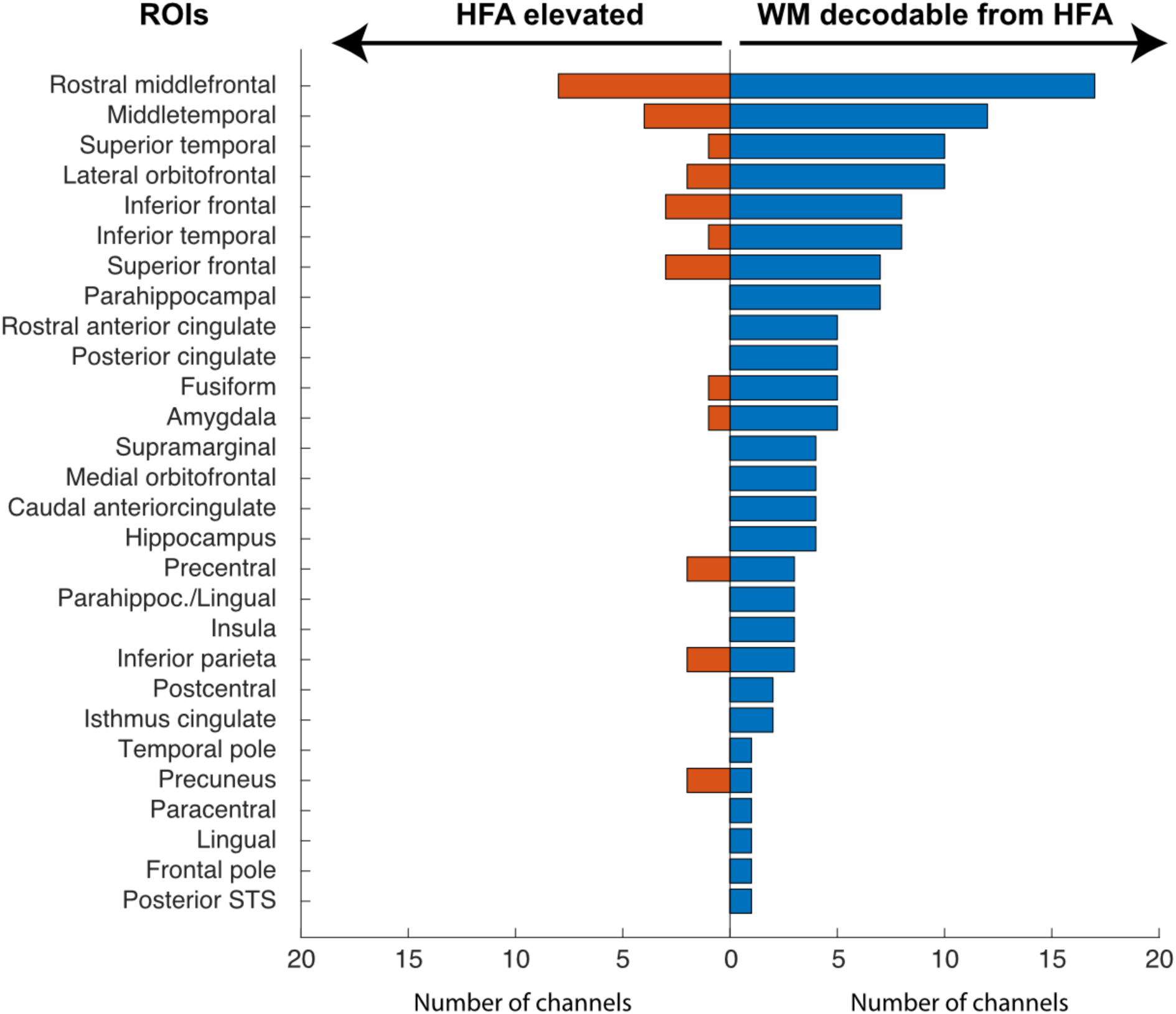
Comparison of the number of channels in each ROI showing elevated HFA (red bars) and significant WM decoding (blue bars) in time-averaged estimates (N=9).

## Discussion

We employed intracranial EEG recordings to gain a more in-depth understanding of the mechanisms of WM. The goal was to test whether the information is maintained by persistent activity or a more dynamic system of activities using MVPA on HFA signal. Our analyses denoted significant increase in decoding accuracy following a response to an irrelevant impulse sound in early AC channels, providing the first direct, local, electrophysiological evidence for activity-silent WM maintenance in the human AC. Our results also reveal dynamic HFA patterns in regions carrying WM content, while only a limited subset of these regions exhibited transient HFA increases without sustained activity.

Impulse-stimulus “perturbation” approaches have previously been utilized to address the challenge of demonstrating activity silent dynamic states as neuronal underpinnings of WM maintenance (Mamashli et al., 2021; Rose et al., 2016; Uluc et al., 2025; Wolff et al., 2015; Wolff et al., 2017b; Wolff et al., 2020). These “perturbation” studies have provided evidence that WM content is maintained in an activity-silent state. That is, information could be decoded by examining content-specific changes to responses to these task-irrelevant impulse stimulus events. However, an essential limitation for these human neuroimaging studies is that the anatomical locus of the WM maintenance can only be indirectly inferred. In line with these findings, we did not find evidence for persistent activity during the WM maintenance period in early AC channels defined by strong responses to the brief impulse sounds (**Figure 2,** left panel). Nevertheless, these same channels showed highly significant increases in decoding accuracy after the impulse sound. These results were consistent at the single participant level and across participants, and they provide robust, spatially resolved evidence for activity silent states enabling WM maintenance, in line with prior suggestions from non-invasive studies in humans (Mamashli et al., 2021; Rose et al., 2016; Wolff et al., 2015; Wolff et al., 2017b; Wolff et al., 2020).

Our findings from the irrelevant impulse sound paradigm provide evidence for activity-silent WM maintenance in the auditory cortex. However, the presence of channels exhibiting elevated HFA is also consistent with recent models proposing the coexistence of persistent neural firing and activity-silent mechanisms as parallel substrates for WM maintenance. A recent interpretation combines these two models for WM maintenance and argues that the underlying mechanism of WM maintenance consists of interactions between high-dimensional, integrative, representations in prefrontal cortex and structured representations in sensory cortex (Buschman, 2021). For example, detailed representations of the sensory stimuli might be stored in sensory cortices. Meanwhile, more abstract aspects of the information could be retained in prefrontal regions (Christophel et al., 2017). The interaction could be explained by the PFC sending a top-down signal to sensory areas, driving oscillations in these areas. The timing of spikes in sensory areas becomes synchronized with these oscillations, and the phase of spiking encodes information about the representations in these areas. Downstream regions can recover this information by combining coherent oscillations and gating input based on the phase of their own local oscillations (Comeaux et al., 2023).

At the whole-brain scale, the dichotomy between neurophysiological recordings in NHPs and human non-invasive studies has elicited different theories on the nature of the WM process, resulting in a reinterpretation of the findings regarding the persistent neuronal firing and alternative activity silent processes during the maintenance period. Representations in both the PFC and sensory cortices can be achieved by persistent spiking alternating between “On” and “Off” states, where during “Off” states, individual neurons lose selectivity for the memorized information and their firing rates return to baseline. However, during these “Off” states, mnemonic information would be still present in the patterns of connections between neuronal groups. This implies that both intermittent neuron spiking and synaptic mechanisms work together to support WM (Panichello et al., 2024). In further support of this theory, attractor dynamics in the PFC, which regulate neural spiking during memory-related periods, has been shown to interact with activity-silent mechanisms in the same region. This interaction enables memory reactivations, which strengthen serial biases in WM (Barbosa et al., 2020). The present results are closely consistent with this dynamic maintenance as they show that content-specific information can be decoded from multiple brain regions, some of which are frontal and auditory regions and hippocampus, with and without elevated HFA.

HFA has been proposed as a proxy of population-level neuronal firing (i.e., MUA) in the immediate vicinity of intracranial depth electrode contacts (Leonard et al., 2024) (for animal studies; Ray et al., 2008; Ray and Maunsell, 2011; but also see; Leszczynski et al., 2020). Persistently elevated MUA patterns in PFC have, in turn, been proposed as a critical component for stable WM maintenance (Fuster et al., 2000; Mendoza-Halliday et al., 2014; Vergara et al., 2016; Constantinidis et al., 2018). Although the persistent firing model of WM does not assume stationary activity at the level of individual units, it predicts that a consistent subset of the population remains engaged throughout the delay period (Constantinidis et al., 2018). At the population level in human sEEG, such patterns would be expected to be manifested as elevated HFA (Constantinidis et al., 2018). Contrary to this prediction, we did not observe sustained increases in mean HFA during WM maintenance across most regions, despite robust decoding of WM content. This pattern is more consistent with a dynamic coding framework, in which WM representations are maintained through time-varying population activity rather than persistent increases in firing rate.

We observed periodical suppression throughout the maintenance period in some of the temporal channels. This is consistent with reports of several fMRI studies (Linke et al., 2011; Ahveninen et al., 2023; Deutsch et al., 2023), as well as with recent multi-site neurophysiological studies that reported suppressed firing in certain brain areas during maintenance (Dotson et al., 2018; Miller et al., 2018). Linke and colleagues have shown that instead of a robust content-specific activation, suppression in fMRI activity in sensory areas enable WM maintenance and dependent on the rehearsal strategy of the participant (Linke et al., 2011). They argue that the suppression, which points to synaptic facilitation rather than persistent firing (Barak & Tsodyks, 2014), might be a natural gatekeeping mechanism against distractors.

In addition to its well-documented role in episodic memory (Burgess et al., 2002; Moscovitch et al., 2016), a growing set of studies provide evidence that HC could play a critical role in WM processing as well (Cabeza et al., 2002; Ezzyat and Olson, 2008; Axmacher et al., 2009; Leszczynski, 2011; Borders et al., 2022). Successful WM performance was shown to be correlated with increased gamma power in HC (Burgess & Ali, 2002; Fell et al., 2001). On the other hand, memory load in successfully remembered trials was correlated with a progressive HC suppression (Stretton et al., 2012). However, HC’s role in WM processing has also been challenged by other earlier (Baddeley et al., 2011) and more recent (Slotnick, 2023) studies. Compatible with research showing HC to be an integral part of WM, our results also show involvement of hippocampus in WM maintenance (**Figure 6**). Our decoding analysis showed that WM information could be found in the HFA from channels recording from HC without the presence of elevated HFA. Borders and colleagues (2021) suggest that HC plays the precise role of supporting complex high-precision binding in WM. Our study design does not allow us to investigate the precision of auditory WM, but our analyses show that HC is involved in maintenance of complex sounds. In combination with the findings in other AC and frontal areas, our results point to a temporally dynamic, spatially distributed network spanning sensory, frontal, and hippocampal regions that is supported by activity silent processes as the neural correlate of WM maintenance.

It should be noted that the sEEG method has limited coverage of the brain. Hence, the activity of other regions contributing to WM maintenance cannot be ruled out. Furthermore, our HFA recordings reflect correlates but not direct measures of underlying neuronal activity (Leszczynski et al., 2020) and, as such, HFA does not allow us to rule out the role of more focal but persistent firing patterns in WM.

In conclusion, our study shows local electrophysiological evidence for the activity silent processes as neural correlates of WM maintenance in early AC. This finding is supported in whole brain level by evidence that neuronal processes underlying WM maintenance are highly dynamic processes that are spread across sensory and frontoparietal areas. These results strongly support the hypothesis that WM maintenance can be supported by a highly dynamic mechanism that is a combination of activity silent processes and population-level neuronal activity.

## Methods

### Participants

We collected sEEG recordings in thirteen participants (n=13, 6 women; age = 33.4 ± 10.9 - ages are jittered for deidentification purposes) who were implanted mainly in frontal and temporal brain regions at Massachusetts General Hospital and Brigham and Women’s Hospital, following the recommendation of a multidisciplinary board. Electrode implantation numbers and locations were determined solely by clinical indications. The participants had varying types of epilepsy with pharmaco-resistant complex partial seizures. Data of one participant was excluded due to insufficient number of trials and 3 participants were excluded due to chance level performance on the WM task. This resulted in 9 analyzed datasets. The data was collected while participants were undergoing invasive monitoring for epilepsy surgical evaluation. All participants provided written, fully informed consent. Participants were informed that participation in the tests would not alter their clinical treatment and that they could withdraw at any time without jeopardizing their clinical care. The study design, protocol, and consent form were approved by the Mass General Brigham Institutional Review Board.

### Stimuli and Procedure

Participants performed a retro-cue WM task with simultaneous recordings of behavior and sEEG from both cortical and subcortical brain structures. In our auditory WM experiment, we employed broadband sound patterns modulated across time (ripple velocity, ω cycles/s) and frequency (Ω cycles/octave) that are called ripple sounds as memory items. Ripple sounds were chosen as memoranda as they have speech-like patterns spectrotemporally (Shamma, 2001) but do not have conceptual properties (Fig. 1a). There were four 750-ms ripple sounds with different ripple velocities that were modified based on our previous studies (Mamashli et al., 2021; Ahveninen et al., 2023). To make the task feasible for our participants during their clinical stay, we determined the ripple velocities based on the largest just-noticeable-difference observed among the participants in previous studies (Mamashli et al., 2021; Ahveninen et al., 2023). The ripple velocities for the four item classes were 9.5, 18.2, 34.9, and 66.8 cycles/s. Two additional velocities were used for probes, including 5.0 and 128 cycles/s. The task was presented via Presentation software (Neurobehavioral Systems, Inc, USA). Sound stimuli were delivered by loudspeakers (JBL Wireless Go 2, Harman I.I.Inc, USA or Cyber Acoustics Sound Bar USB Speaker, Cyber Acoustics inc., USA) externally connected to the task presentation computer.

The retro-cue WM paradigm, which is well established in the field (Christophel et al., 2012; Christophel and Haynes, 2014; Uluç et al., 2018; Ahveninen et al., 2023), aims to dissociate perception and attention from memory maintenance (**Fig. 8b**). It is essentially a modification of a delayed match to sample paradigm. In our WM task, we sequentially presented two 750-ms ripple sounds with a 250-ms interstimulus interval (ISI). The second ripple sound was followed by a retro-cue indicating the item to remember. The visual retro-cue consisted of a visual “memorize1” or “memorize2” cue after another 250-ms ISI. After a 3-s maintenance period, in the middle of which an unrelated sound stimulus (50-ms white noise)—i.e., an “impulse stimulus” (Wolff, Jochim, Akyurek, & Stokes, 2017a) —was presented, a ripple sound probe was presented with a same/different response scheme. Participant gave a self-paced response by pressing A (same) or S (different) button presses on the keyboard, which were labeled respectively as “1” and “2”, indicating if the test probe is the same as the memorized item or not. After each trial, a 1-s inter trial interval was registered (**Fig. 8b**).

**Figure 8.**
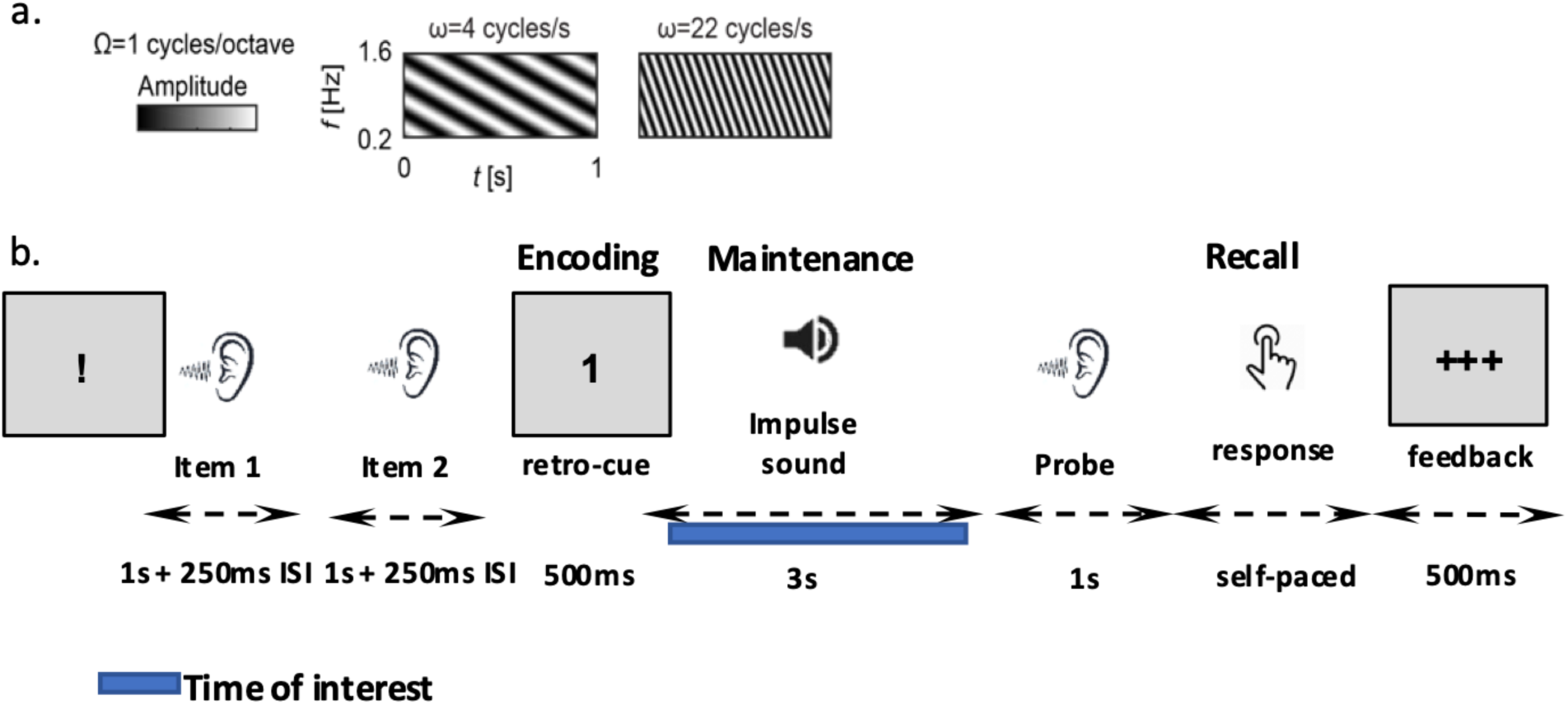
Diagram of the stimuli and paradigm. **(a)** Visualization of the spectrotemporal pattern of ripple sound stimuli. **(b)** Time course of the WM task. The blue bar represents the maintenance period during which sEEG data were analyzed. The presentation of ripple sounds is denoted by waves to the ears and the impulse sound is denoted by a loudspeaker symbol. Participants were instructed to indicate if the probe matched the item indicated by the retro-cue via a button press in a self-paced manner and were provided with visual feedback.

Each run consisted of 24 trials, 12 of which were match and the remaining non-match. The non-match trials were divided into two parts with half the trials complete non-match trials where the probe was a different ripple sound from the two items that are presented at the beginning of the trial. The other half of the non-match trials designated the probe as the uncued item. The memory items were pseudo-randomly chosen from four different classes with different temporal frequencies. To eliminate categorical encoding, participants were naïve to the number of items to memorize. The task was planned to be 120 trials in total if completed with 5 runs. The number of trials/runs completed varied across participants.

### Data Acquisition

sEEG was recorded from a montage of 8-18 unilaterally or bilaterally implanted depth electrodes depending on the clinically indicated procedure per participant. Intracranial signals were recorded using a recording system with a sampling rate of 2000 Hz (Neural Signal Processors, Blackrock Microsystems, US) or with a sampling rate of 1024 Hz (Natus Quantum). At the time of acquisition, sEEG recordings were referenced to an EEG electrode that was placed on the head (either mastoid, cervical vertebrae 2, Cz or a sEEG contact that, based on imaging localization, was located in the skull and outside the brain). Intracranial electrodes were localized by an established volumetric image co-registration procedure that is used for clinical application. First, the pre-operation T1-weighted MRI images were aligned to post-operation CT or MRI images that show electrode locations using Freesurfer scripts (http://surfer.mnr.mgh.harvard.edu). Electrode coordinates were manually determined from the post operation implant images. The electrodes were then mapped to standard cortical parcels/regions using FreeSurfer, assigning the RAS coordinates of each contact location to their closest cortical surface label (Dykstra et al., 2012; Soper et al., 2023). For this purpose, we generated a volumetric version of the FreeSurfer cortical parcellation. Each participant’s original cortical parcellation was first transformed into volumetric space using FreeSurfer’s mri_label2vol, to ensure that the channels deeper in the cortical ribbon are also included in the appropriate labels. Two complementary projections were used: A fractional-depth projection, obtaining cortical labels across the full thickness of the ribbon and an absolute-depth projection, extending labels further into the underlying white matter to ensure robustness in deeper locations. These two projections were combined to produce a more complete volumetric label set. The resulting volumetric label map provided a continuous 3D representation of cortical parcellations. This enabled direct assignment of intracranial electrode locations (or other volumetric data) to the nearest anatomical label. The analyses were concentrated on the cortical parcels of the Freesurfer Desikan atlas, as well as the hippocampus (HC).

### Preprocessing

sEEG data was preprocessed with custom MATLAB scripts using the Fieldtrip toolbox (Oostenveld et al., 2011). The data was divided into single-trial epochs starting from 300 ms before to 3 s after the retro-cue onset. Noisy channels as well as channels with excessive epileptiform activity and bad epochs were identified and rejected based on visual evaluation. The data was detrended with a baseline window of −300 ms to 0 ms and filtered with a 300 Hz low pass and a 0.1 Hz high pass filter sequentially using *ft_preprocessing*. Notch filtering was used to remove 60 Hz line noise and its harmonics from the data. Whole trial datasets were detrended and both European data format (EDF) and Blackrock format datasets were downsampled to 1 kHz. Data was then re-referenced to a common average of all remaining electrodes. Afterwards, using an automatic interictal discharge detection algorithm (Janca et al., 2015), we removed the channels that had 6.5 discharges/minute as pathological channels. In the final analyses, we used Laplacian re-referencing, following prior work that supports its application for local population-level SEEG activity (Li et al., 2018). By suppressing uncorrelated noise in the reference signal through spatial averaging, this approach was presumed to provide favorable SNR characteristics for MVPA analyses. In this case, a channel indicates the recording which can be from individual or re-referenced contacts per electrode. Trials with artifacts were removed from all channels regardless of the channel signal to ensure the analyses were not confounded in any of the channels.

### Main analyses

#### HFA Analysis

To examine the evolution of HFA power during the task trials, Laplacian-referenced sEEG epochs spanning −300 ms and 9 s relative to the onset cue were band-pass filtered between 70-190 Hz, Hilbert transformed, squared, and baseline corrected using the 300-ms pre-cue period. To identify patterns of elevated delay activity, HFA was averaged during the maintenance period while excluding the first 500 ms after both the visual retro-cue and auditory impulse stimulus onsets, which might have contained unrelated event-related activity.

#### Statistical Significance of HFA Results

The statistical significance of HFA changes relative to baseline was evaluated within each participant using a one-sample permutation test based on the *t*-statistic across all trials. To handle multiple comparisons across all participants, we determined the maximum critical *t* threshold across all participants and compared each participant’s *t* values from their regions of interest (ROIs) to this threshold. HFA changes in ROIs with t values exceeding the (between-participant) maximum critical *t* value were deemed statistically significant after multiple comparison correction.

#### Multivariate Pattern Analysis

We decoded the 4 ripple sound memory items that participants had to memorize from the WM maintenance period. Participants were naïve to the number of memory items to discourage conceptualization. The within participant 4-class classification analysis was conducted by employing the SVM implementation from libsvm (Chang & Lin, 2011) and MATLAB/Octave CoSMoMVPA package (Oosterhof, Connolly, & Haxby, 2016). To estimate content-specific patterns of HFA during WM maintenance, Laplacian-referenced sEEG epochs spanning −300 ms and 3 s relative to the onset of the retro-cue were baseline corrected using the 300-ms pre-cue period and band-pass filtered between 70-190 Hz. The numbers of trials per class were balanced utilizing CoSMoMVPA and Fieldtrip toolboxes. To improve the signal-to-noise ratio (SNR), we calculated averages of four trials for each class (using *cosmo_average_samples*). We then performed 20 random iterations using these averaged samples.

A 4-class classification was carried out using a temporal searchlight analysis (Oosterhof et al., 2016) with a 300 ms sliding window in 3 ms steps, conducted for each contact separately. A k-fold cross-validation procedure was employed to classify the maintained WM content. In each fold, the model was trained on k-1/k of the samples and tested on the remaining samples. Decoding accuracies for each searchlight dataset per contact were averaged across folds and iterations. For each participant and contact, this analysis produced a time series with decoding accuracies for each searchlight centroid.

#### Statistical Significance of MVPA Results

The statistical significance of our MVPA results was assessed by employing robust non-parametric permutation tests (Mamashli et al., 2021), which handle multiple comparison problems using a *between-participant maximum-statistic* strategy. In this procedure, we first generated 500 unique permutations of the true item-content labels of the classifier. The temporal searchlight analysis was repeated with these permuted labels to generate a distribution of decoding accuracies for each time point. For each permutation, the time series of decoding accuracies were converted to *z*-values. This was done by comparing each decoding-accuracy value to the respective permutation distribution at the same time point.

Hypothesis testing was implemented using two complementary strategies. The primary analysis aimed to identify sEEG channels with statistically significant decoding over the same maintenance-period time segments that had been used for analyzing HFA power changes. To achieve this, z values representing normalized decoding accuracies were averaged across the maintenance period, while excluding searchlight periods overlapping with the first 500 ms following both the visual retro-cue and the auditory impulse stimulus onsets. This procedure was repeated for 500 permutations.

For each participant and hemisphere, the maximum z value from each permutation was used to construct a participant- and hemisphere-specific initial null distribution. Finally, the maximum z values across all participants and hemispheres were pooled to form the final null distribution. The statistical significance of normalized decoding accuracies for each participant and contact was determined by comparing observed values against this between-participant maximum-statistic null distribution.

Additionally, to explore the temporal dynamics of decoding accuracy, we used a cluster-based analysis based on the same permutation distributions that had been calculated for the main analysis. Continuous clusters with *Z*>2.3 (i.e., *p*<0.01) were then identified in each permutation and the respective cluster sums of *Z-*values were calculated. From each permutation, the largest cluster sum across all conditions was entered to the maximum-statistic null distribution in each hemisphere and participant. Finally, the maximum cluster sums across all participants and hemispheres were pooled to form the final null distribution. The statistical significance of clusters in each participant and contact was determined by comparing observed cluster-sum values against this between-participant maximum-statistic null distribution.

## Acknowledgements

This project was supported by NIH grants R01DC017991, R01DC016765, R01DC016915, P41EB030006, R01-2NS062092-0 and Tiny Blue Dot grant P50MH119467-01A1.

