## Supplementary material for "Human intracranial correlates of dynamic coding in auditory working memory"

### Supplementary Information

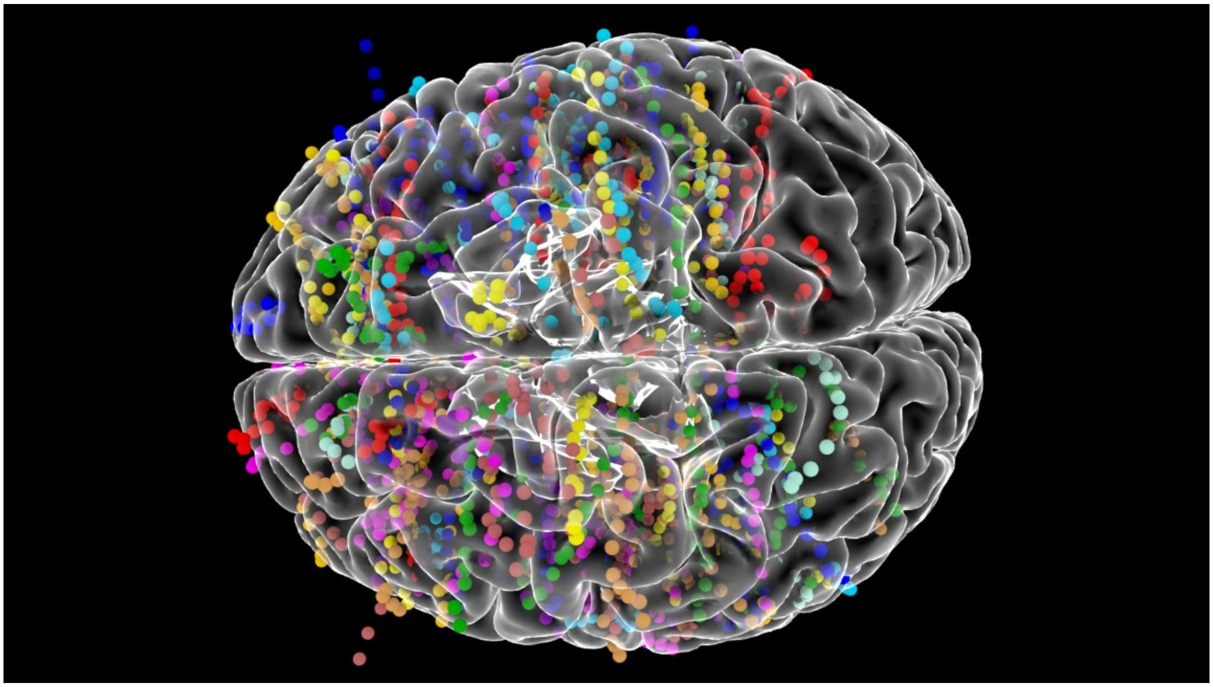

**Figure S1.** Coverage for all contacts across all subjects normalized to Colin27 common space. Contacts of each individual participant have been labeled with a different color.

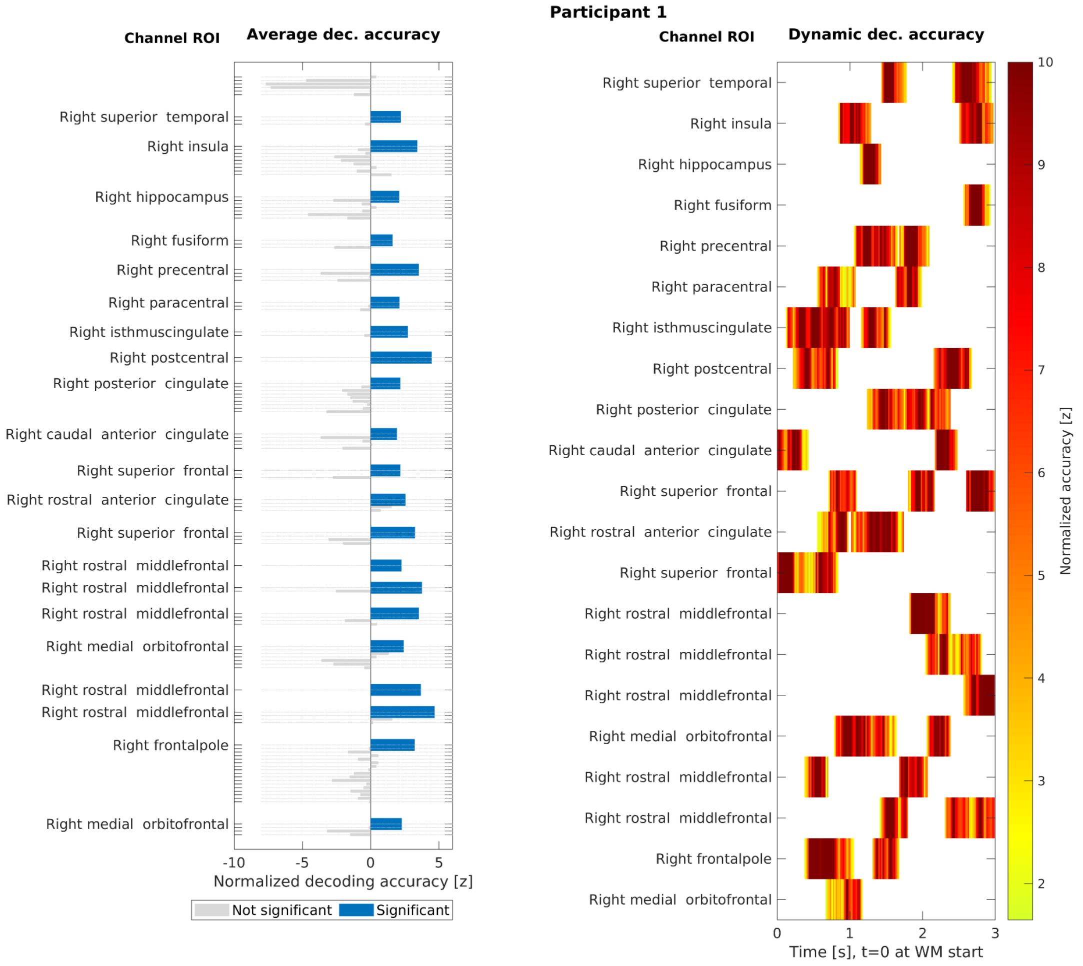


**Figure S2**. Maintenance period 4-class SVM classification results in Participant 1. (Left) Average normalized decoding accuracy in all channels. The channels, where decoding accuracy was, on average, significantly above chance level, have been denoted with blue bars. Statistical significance was determined by a permutation test with a between-subject maximum-statistic null distribution. (Right) MVPA time courses masked to statistically significant clusters, focused on channels also significant in the time-averaged test. The channels are anatomically labeled based on Freesurfer atlas. Statistical significance of the clusters was determined using cluster-based permutation test with a between-subject maximum-statistic null distribution.

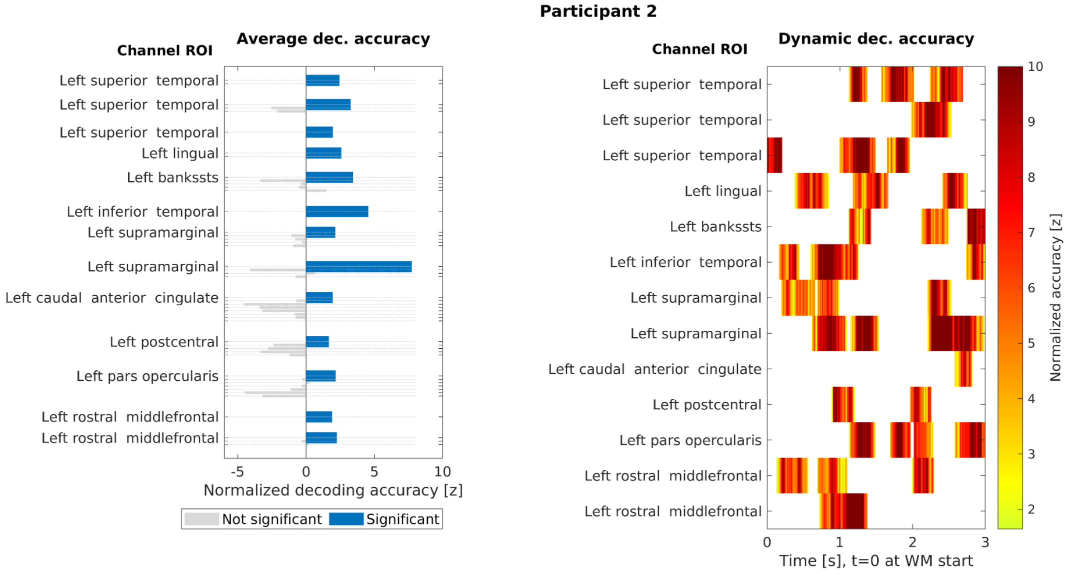


**Figure S3**. Maintenance period 4-class SVM classification results in Participant 2. (Left) Average normalized decoding accuracy in all channels. The channels, where decoding accuracy was, on average, significantly above chance level, have been denoted with blue bars. Statistical significance was determined by a permutation test with a between-subject maximum-statistic null distribution. (Right) MVPA time courses masked to statistically significant clusters, focused on channels also significant in the time-averaged test. The channels are anatomically labeled based on Freesurfer atlas. Statistical significance of the clusters was determined using cluster-based permutation test with a between-subject maximum-statistic null distribution.

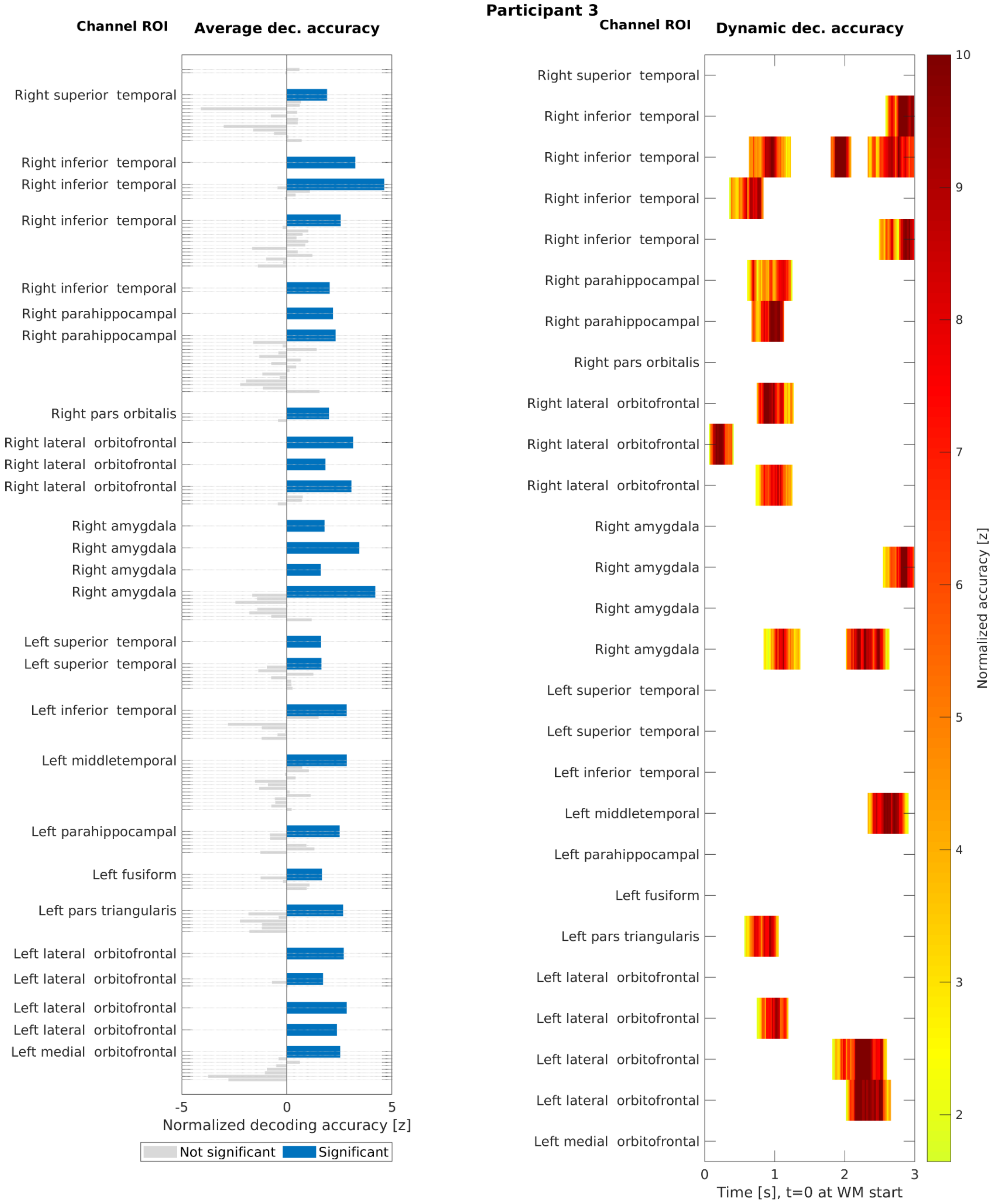


**Figure S4**. Maintenance period 4-class SVM classification results in Participant 3. (Left) Average normalized decoding accuracy in all channels. The channels, where decoding accuracy was, on average, significantly above chance level, have been denoted with blue bars. Statistical significance was determined by a permutation test with a between-subject maximum-statistic null distribution. (Right) MVPA time courses masked to statistically significant clusters, focused on channels also significant in the time-averaged test. The channels are anatomically labeled based on Freesurfer atlas. Statistical significance of the clusters was determined using cluster-based permutation test with a between-subject maximum-statistic null distribution.

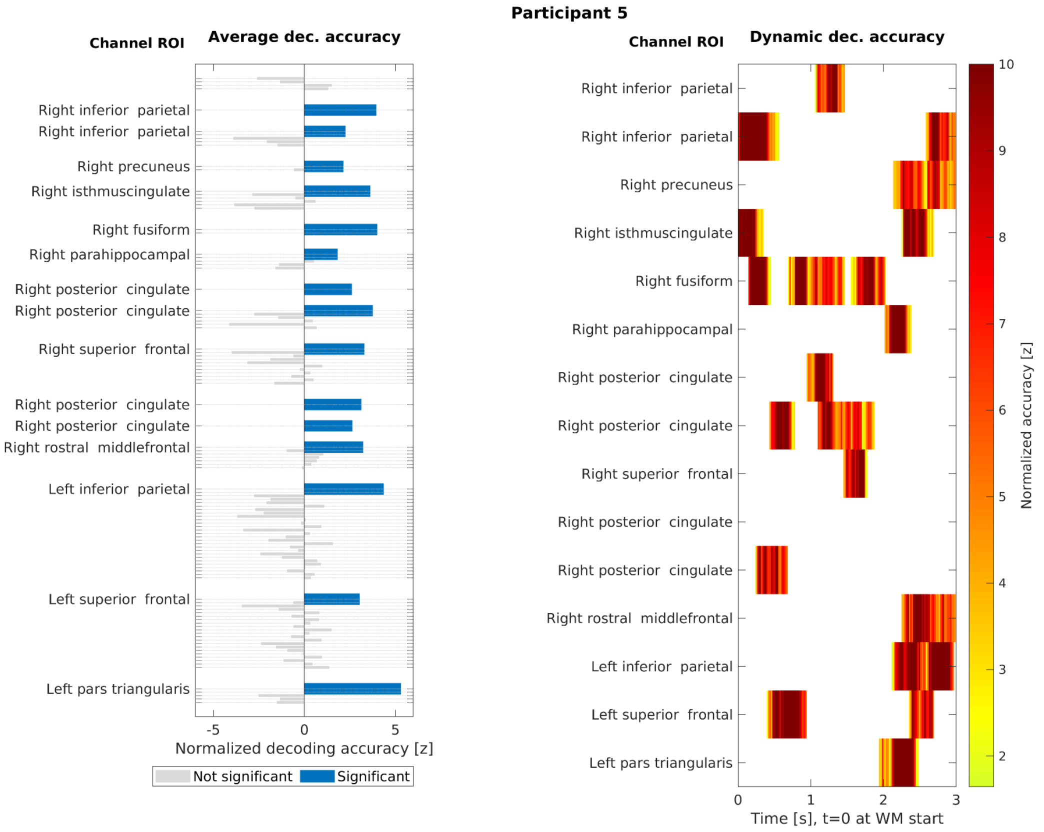


**Figure S5**. Maintenance period 4-class SVM classification results in Participant 5. (Left) Average normalized decoding accuracy in all channels. The channels, where decoding accuracy was, on average, significantly above chance level, have been denoted with blue bars. Statistical significance was determined by a permutation test with a between-subject maximum-statistic null distribution. (Right) MVPA time courses masked to statistically significant clusters, focused on channels also significant in the time-averaged test. The channels are anatomically labeled based on Freesurfer atlas. Statistical significance of the clusters was determined using cluster-based permutation test with a between-subject maximum-statistic null distribution.

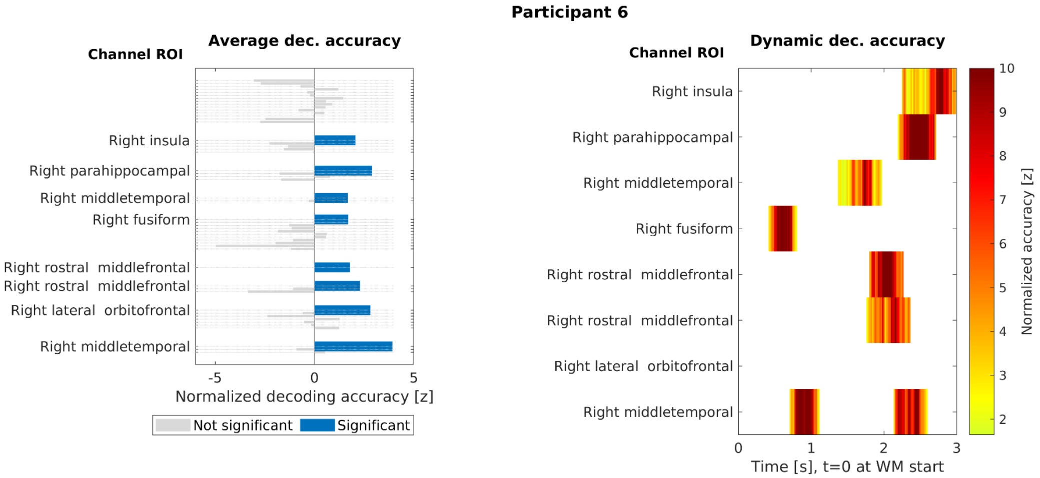


**Figure S6**. Maintenance period 4-class SVM classification results in Participant 6. (Left) Average normalized decoding accuracy in all channels. The channels, where decoding accuracy was, on average, significantly above chance level, have been denoted with blue bars. Statistical significance was determined by a permutation test with a between-subject maximum-statistic null distribution. (Right) MVPA time courses masked to statistically significant clusters, focused on channels also significant in the time-averaged test. The channels are anatomically labeled based on Freesurfer atlas. Statistical significance of the clusters was determined using cluster-based permutation test with a between-subject maximum-statistic null distribution.

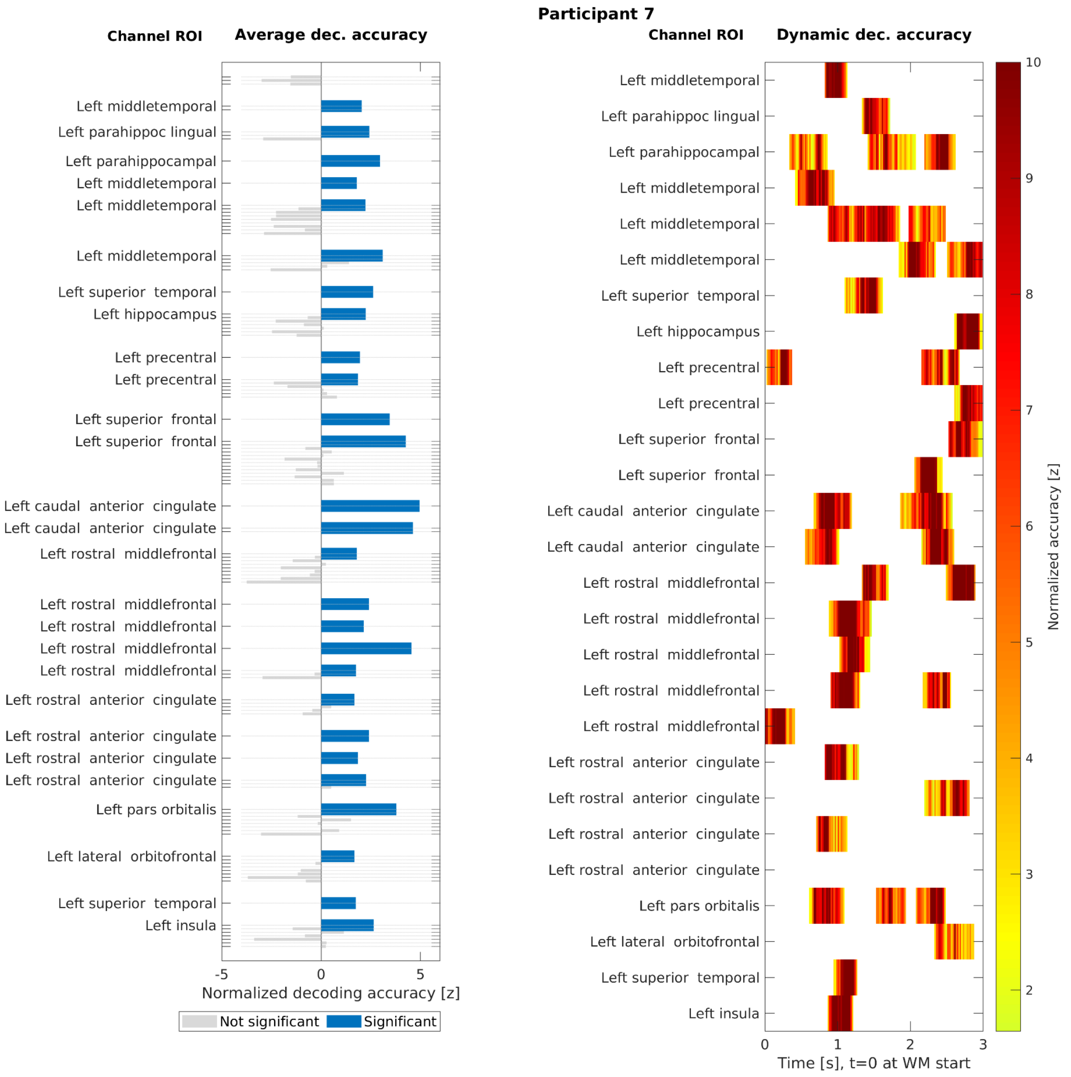


**Figure S7**. Maintenance period 4-class SVM classification results in Participant 7. (Left) Average normalized decoding accuracy in all channels. The channels, where decoding accuracy was, on average, significantly above chance level, have been denoted with blue bars. Statistical significance was determined by a permutation test with a between-subject maximum-statistic null distribution. (Right) MVPA time courses masked to statistically significant clusters, focused on channels also significant in the time-averaged test. The channels are anatomically labeled based on Freesurfer atlas. Statistical significance of the clusters was determined using cluster-based permutation test with a between-subject maximum-statistic null distribution.

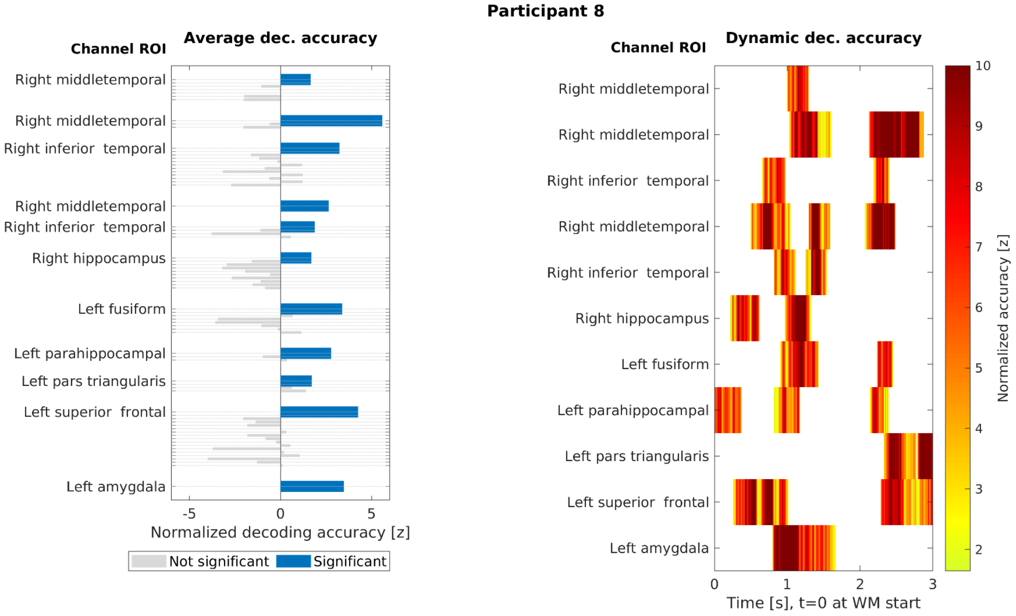


**Figure S8**. Maintenance period 4-class SVM classification results in Participant 8. (Left) Average normalized decoding accuracy in all channels. The channels, where decoding accuracy was, on average, significantly above chance level, have been denoted with blue bars. Statistical significance was determined by a permutation test with a between-subject maximum-statistic null distribution. (Right) MVPA time courses masked to statistically significant clusters, focused on channels also significant in the time-averaged test. The channels are anatomically labeled based on Freesurfer atlas. Statistical significance of the clusters was determined using cluster-based permutation test with a between-subject maximum-statistic null distribution.

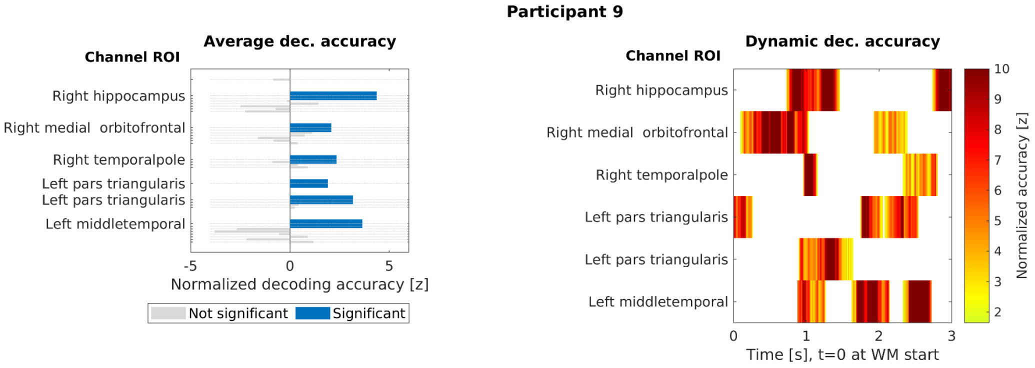


**Figure S9**. Maintenance period 4-class SVM classification results in Participant 9. (Left) Average normalized decoding accuracy in all channels. The channels, where decoding accuracy was, on average, significantly above chance level, have been denoted with blue bars. Statistical significance was determined by a permutation test with a between-subject maximum-statistic null distribution. (Right) MVPA time courses masked to statistically significant clusters, focused on channels also significant in the time-averaged test. The channels are anatomically labeled based on Freesurfer atlas. Statistical significance of the clusters was determined using cluster-based permutation test with a between-subject maximum-statistic null distribution.

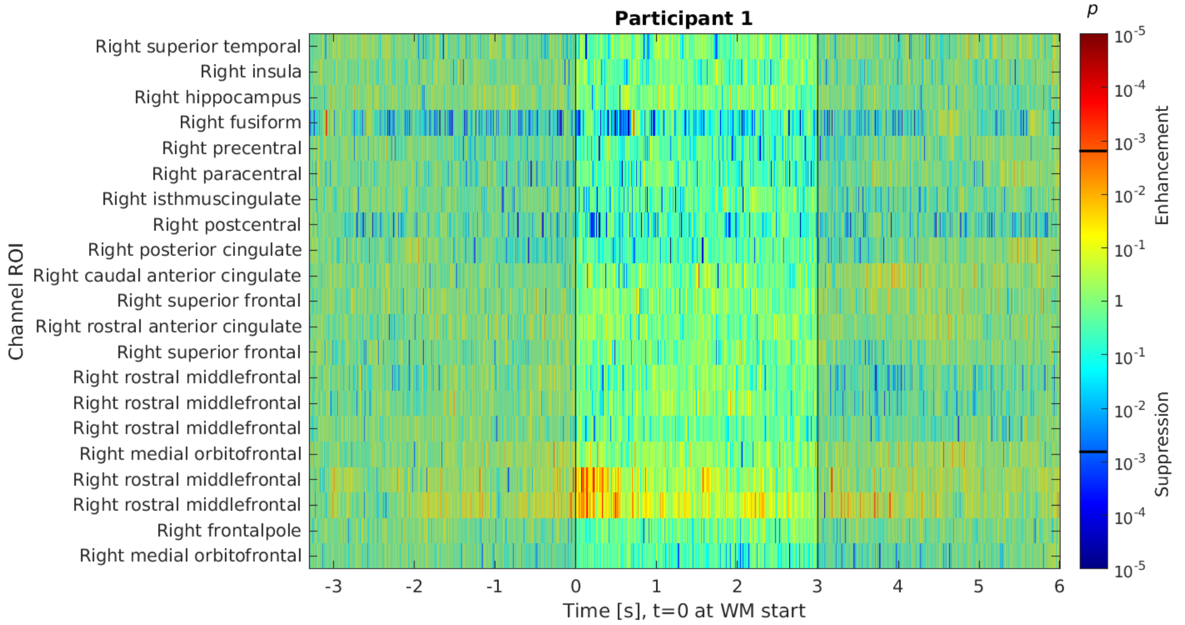


**Figure S10**. Whole-trial averaged HFA analysis from the channels that carried information of the WM content in Participant 1. The time periods before and after WM maintenance period are shaded. The horizontal black lines in the color bar refer to the critical p value determined based on FDR.

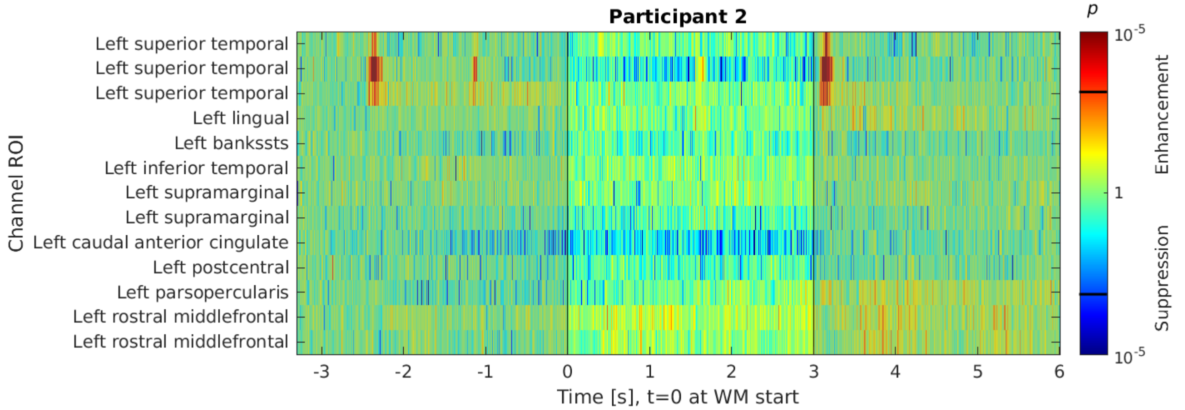


**Figure S11**. Whole-trial averaged HFA analysis from the channels that carried information of the WM content in Participant 2. The time periods before and after WM maintenance period are shaded. The horizontal black lines in the color bar refer to the critical p value determined based on FDR.

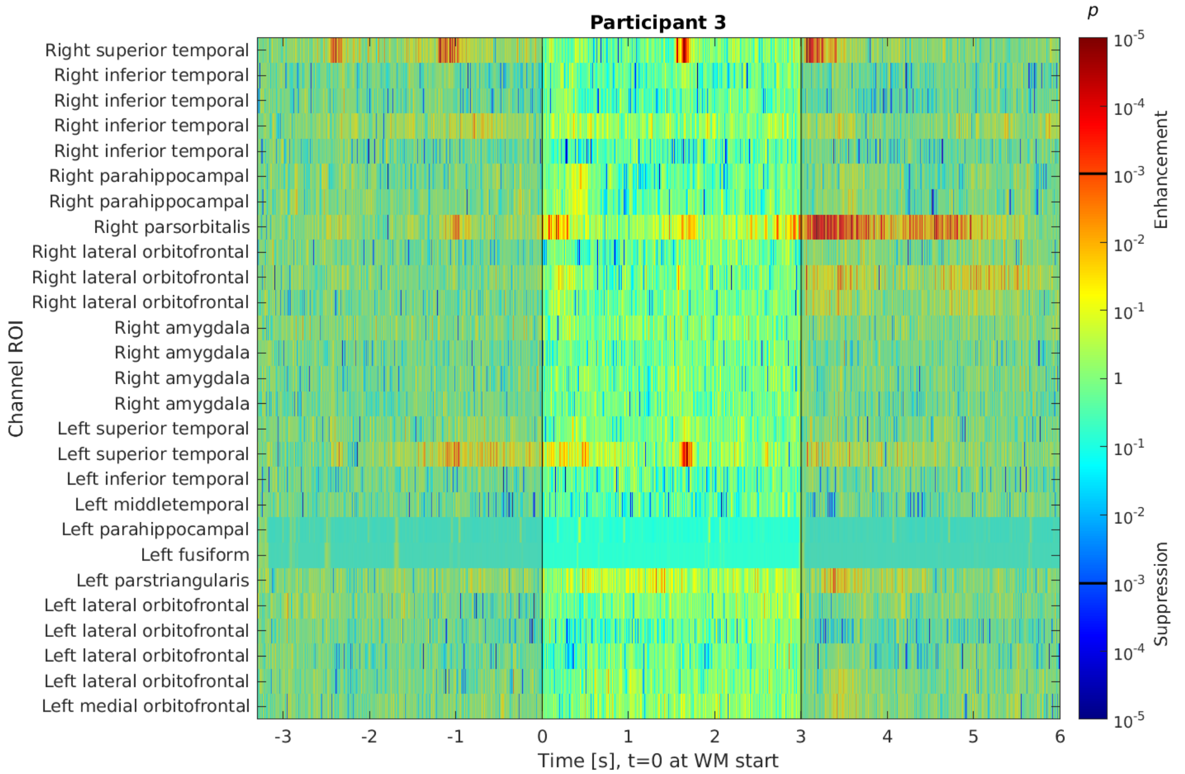


**Figure S12**. Whole-trial averaged HFA analysis from the channels that carried information of the WM content in Participant 3. The time periods before and after WM maintenance period are shaded. The horizontal black lines in the color bar refer to the critical p value determined based on FDR.

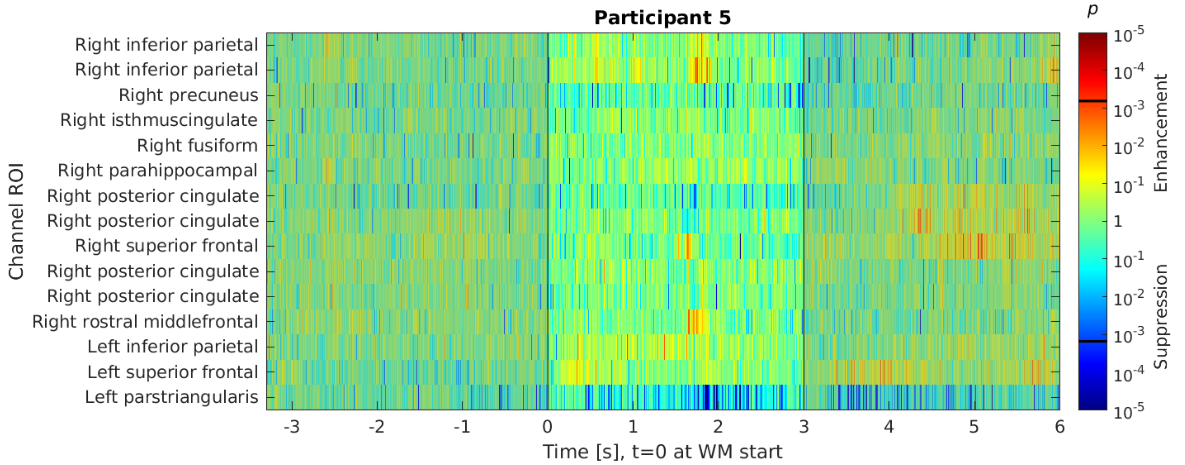


**Figure S13**. Whole-trial averaged HFA analysis from the channels that carried information of the WM content in Participant 5. The time periods before and after WM maintenance period are shaded. The horizontal black lines in the color bar refer to the critical p value determined based on FDR.

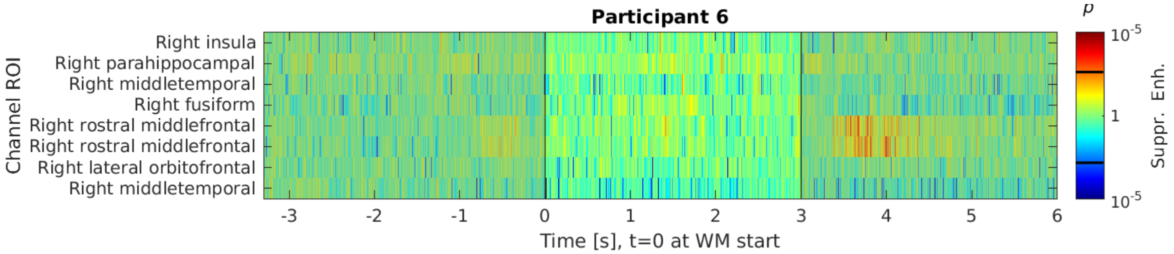


**Figure S14**. Whole-trial averaged HFA analysis from the channels that carried information of the WM content in Participant 6. The time periods before and after WM maintenance period are shaded. The horizontal black lines in the color bar refer to the critical p value determined based on FDR.

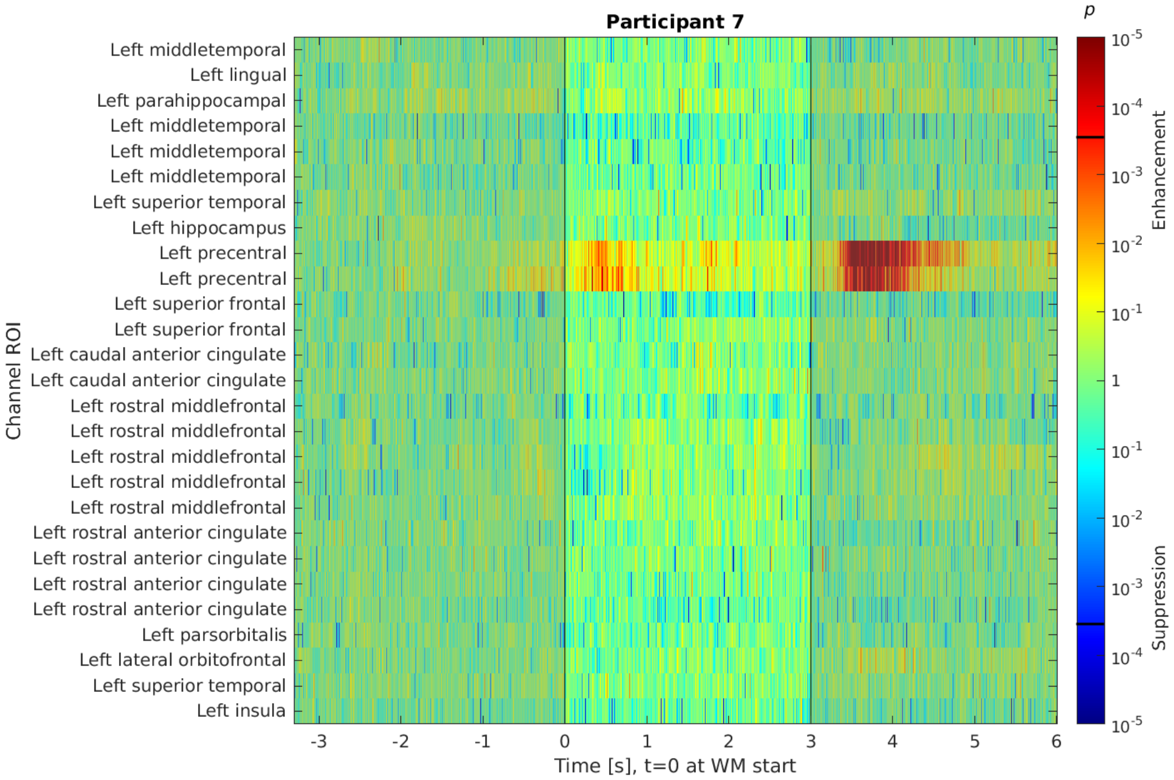


**Figure S15**. Whole-trial averaged HFA analysis from the channels that carried information of the WM content in Participant 7. The time periods before and after WM maintenance period are shaded. The horizontal black lines in the color bar refer to the critical p value determined based on FDR.

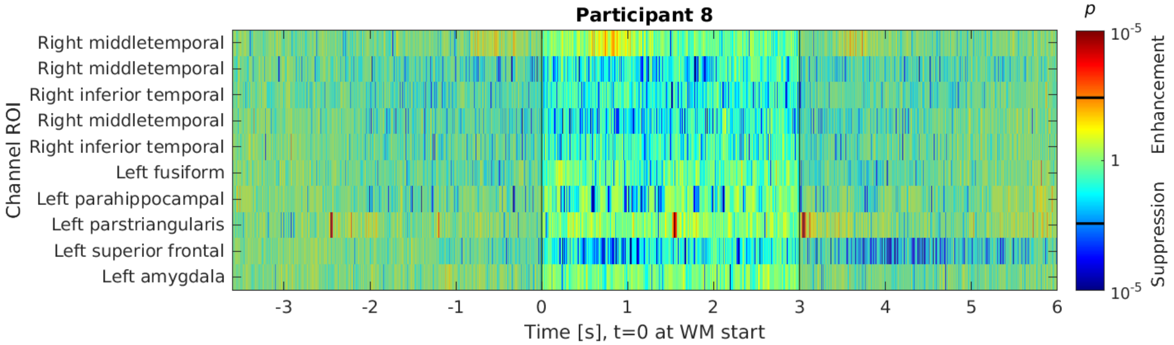


**Figure S16**. Whole-trial averaged HFA analysis from the channels that carried information of the WM content in Participant 8. The time periods before and after WM maintenance period are shaded. The horizontal black lines in the color bar refer to the critical p value determined based on FDR.

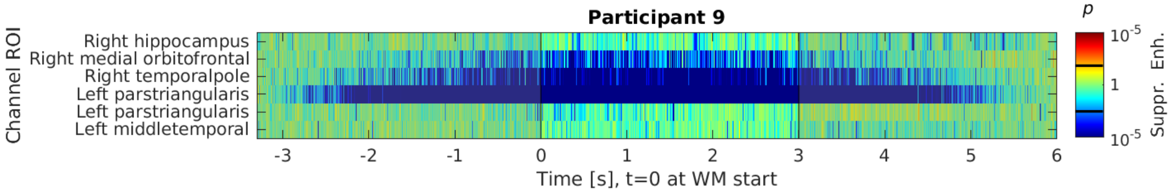


**Figure S17**. Whole-trial averaged HFA analysis from the channels that carried information of the WM content in Participant 9. The time periods before and after WM maintenance period are shaded. The horizontal black lines in the color bar refer to the critical p value determined based on FDR.

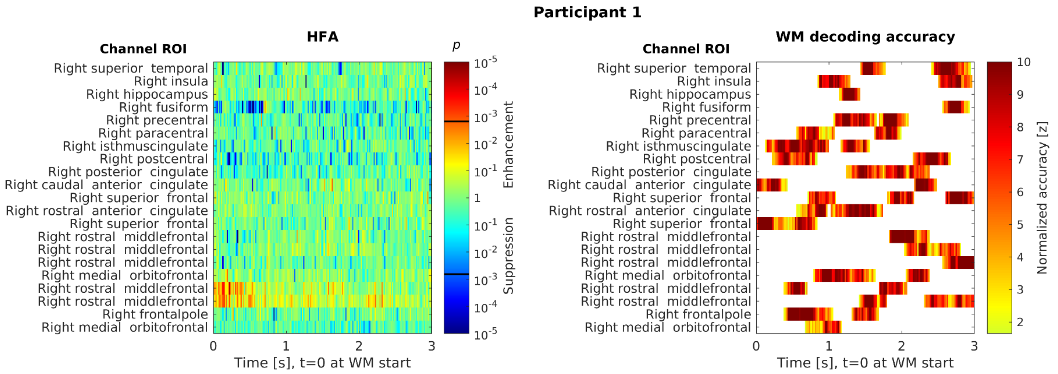


**Figure S18**. HFA and MVPA results for WM maintenance period in Participant 1 in channels carrying information of WM content. The channels have been anatomically labeled based on the Freesurfer atlas. (Left) HFA time course for the WM maintenance period. The horizontal black lines in the HFA color bar refer to the critical p value determined based on FDR. (Right) MVPA results for WM maintenance period. The decoding results have been thresholded based on a cluster-based permutations test with a between-subject maximum-statistic null distribution.

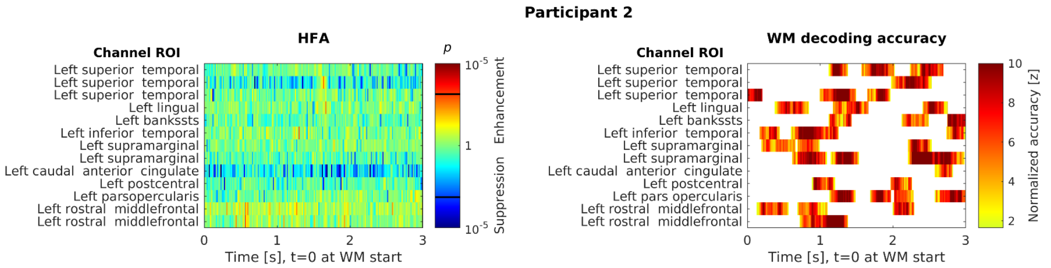


**Figure S19**. HFA and MVPA results for WM maintenance period in Participant 2 in channels carrying information of WM content. The channels have been anatomically labeled based on the Freesurfer atlas. (Left) HFA time course for the WM maintenance period. The horizontal black lines in the HFA color bar refer to the critical p value determined based on FDR. (Right) MVPA results for WM maintenance period. The decoding results have been thresholded based on a cluster-based permutations test with a between-subject maximum-statistic null distribution.

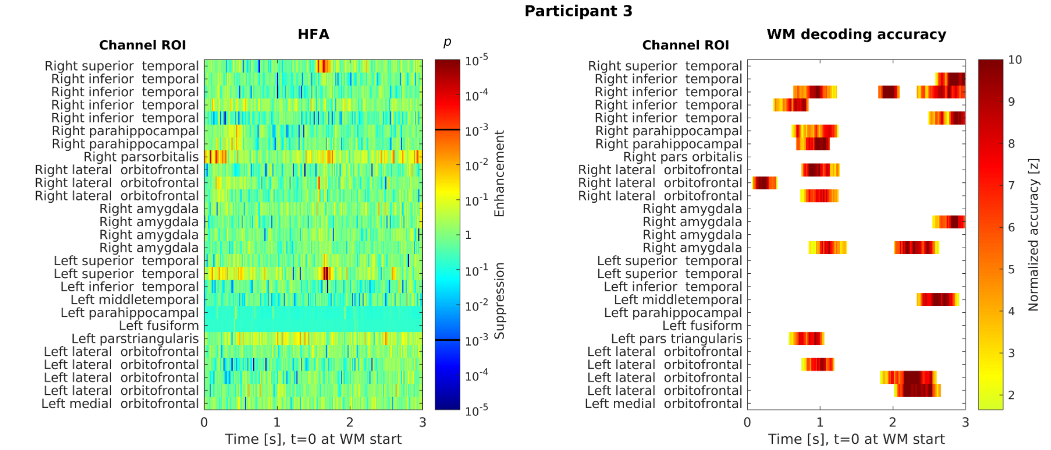


**Figure S20**. HFA and MVPA results for WM maintenance period in Participant 3 in channels carrying information of WM content. The channels have been anatomically labeled based on the Freesurfer atlas. (Left) HFA time course for the WM maintenance period. The horizontal black lines in the HFA color bar refer to the critical p value determined based on FDR. (Right) MVPA results for WM maintenance period. The decoding results have been thresholded based on a cluster-based permutations test with a between-subject maximum-statistic null distribution.

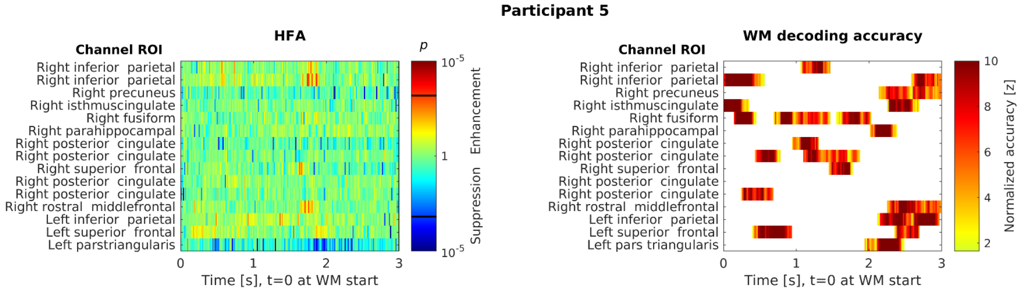


**Figure S21**. HFA and MVPA results for WM maintenance period in Participant 5 in channels carrying information of WM content. The channels have been anatomically labeled based on the Freesurfer atlas. (Left) HFA time course for the WM maintenance period. The horizontal black lines in the HFA color bar refer to the critical p value determined based on FDR. (Right) MVPA results for WM maintenance period. The decoding results have been thresholded based on a cluster-based permutations test with a between-subject maximum-statistic null distribution.

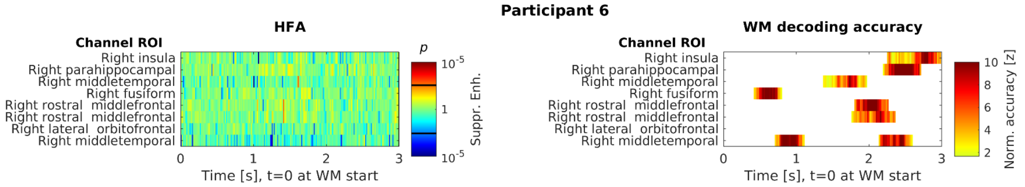


**Figure S22**. HFA and MVPA results for WM maintenance period in Participant 6 in channels carrying information of WM content. The channels have been anatomically labeled based on the Freesurfer atlas. (Left) HFA time course for the WM maintenance period. The horizontal black lines in the HFA color bar refer to the critical p value determined based on FDR. (Right) MVPA results for WM maintenance period. The decoding results have been thresholded based on a cluster-based permutations test with a between-subject maximum-statistic null distribution.

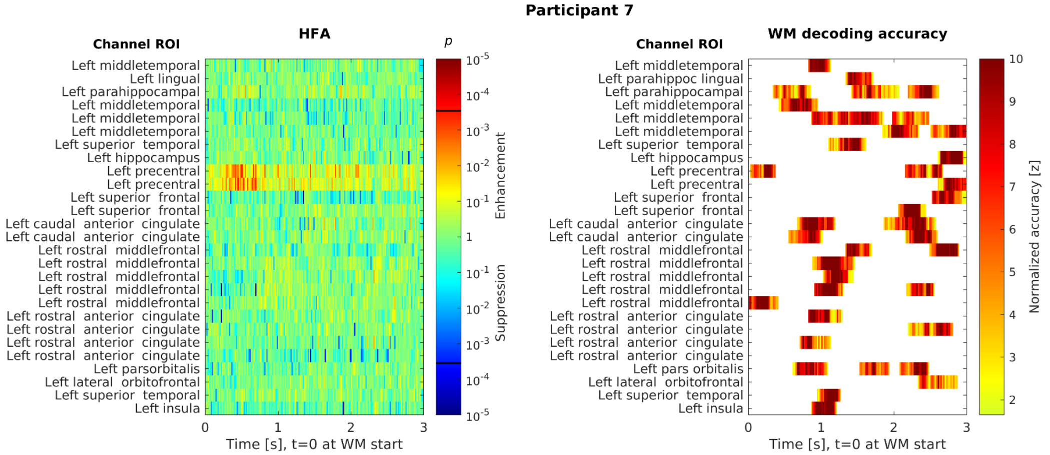


**Figure S23**. HFA and MVPA results for WM maintenance period in Participant 7 in channels carrying information of WM content. The channels have been anatomically labeled based on the Freesurfer atlas. (Left) HFA time course for the WM maintenance period. The horizontal black lines in the HFA color bar refer to the critical p value determined based on FDR. (Right) MVPA results for WM maintenance period. The decoding results have been thresholded based on a cluster-based permutations test with a between-subject maximum-statistic null distribution.

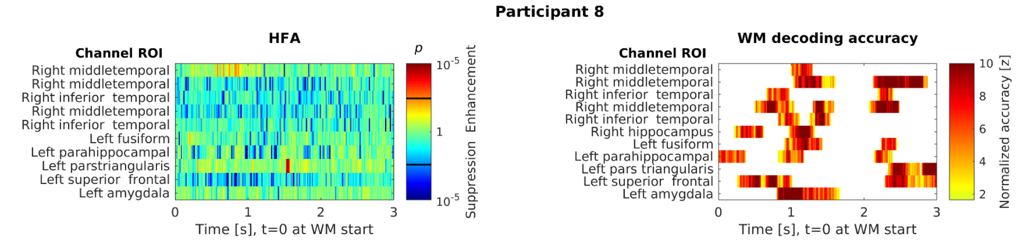


**Figure S24**. HFA and MVPA results for WM maintenance period in Participant 8 in channels carrying information of WM content. The channels have been anatomically labeled based on the Freesurfer atlas. (Left) HFA time course for the WM maintenance period. The horizontal black lines in the HFA color bar refer to the critical p value determined based on FDR. (Right) MVPA results for WM maintenance period. The decoding results have been thresholded based on a cluster-based permutations test with a between-subject maximum-statistic null distribution.

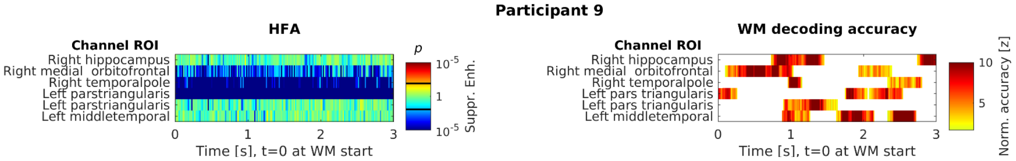


**Figure S25**. HFA and MVPA results for WM maintenance period in Participant 9 in channels carrying information of WM content. The channels have been anatomically labeled based on the Freesurfer atlas. (Left) HFA time course for the WM maintenance period. The horizontal black lines in the HFA color bar refer to the critical p value determined based on FDR. (Right) MVPA results for WM maintenance period. The decoding results have been thresholded based on a cluster-based permutations test with a between-subject maximum-statistic null distribution.

**Table S1**. Implantation sites per participant. All electrode implantation plans were dictated by clinical necessity only. The number of contacts denotes the total number of contacts that are recorded on the implanted hemisphere. Participants were denoted according to the names given in the main analysis.

| Participant Nr | Implantation Site | | | |
|---|---|---|---|---|
| | Left Hemisphere | Nr of contacts | Right Hemisphere | Nr of contacts |
| **Participant 1** | No | - | Yes | 176 |
| **Participant 2** | Yes | 112 | No | - |
| **Participant 3** | Yes | 134 | Yes | 122 |
| **Participant 4** | No | - | Yes | 158 |
| **Participant 5** | Yes | 112 | Yes | 98 |
| **Participant 6** | No | - | Yes | 126 |
| **Participant 7** | Yes | 184 | No | - |
| **Participant 8** | Yes | 62 | Yes | 62 |
| **Participant 9** | Yes | 30 | Yes | 30 |
| **Participant 10** | Yes | 62 | Yes | 20 |
| **Participant 11** | Yes | 146 | No | - |
| **Participant 12** | No | - | Yes | 112 |
| **Participant 13** | Yes | 142 | Yes | 24 |

**Table S2**. Clinical information per participant.

| | Handedness | Previous resection | Etiology, region of noted area of intracranial seizure activity |
|---|---|---|---|
| **Participant 1** | Right | Yes, right anterior frontal and temporal lobe | Unknown, Right Fronto-temporal lobes |
| **Participant 2** | Right | No | Unknown, left insula |
| **Participant 3** | Right | No | Unknown, left mesial temporal lobe and temporal pole |
| **Participant 4** | Right | No | Unknown, right mesial temporal lobe and temporal pole |
| **Participant 5** | No | Yes, right frontal lobe | Trauma, right frontal lobe |
| **Participant 6** | Left | No | Unknown, right mesial temporal lobe |
| **Participant 7** | Right | No | Unknown, left superficial frontal lobe |
| **Participant 8** | Both | No | Unknown, bilateral mesial temporal lobe |
| **Participant 9** | Both | No | Unknown, bilateral mesial temporal lobe |
| **Participant 10** | Left | Yes, left temporal lobe | Unknown, left mesial temporal lobe |
| **Participant 11** | Right | Yes, left temporal lobe | Tumor, left mesial temporal lobe |
| **Participant 12** | Right | No | Unknown, right mesial temporal lobe |
| **Participant 13** | Right | No | Unknown, results unclear |
